# Combined sulfur deficiency and water deficit trigger synergistic redox adjustments through coordinated transcript-protein regulation in pea

**DOI:** 10.1101/2024.03.13.582463

**Authors:** Titouan Bonnot, Charlotte Henriet, Delphine Aimé, Jonathan Kreplak, Morgane Térézol, Nadia Rossin, Thierry Balliau, Cécile Blanchard, Olivier Lamotte, Alain Ourry, Michel Zivy, Vanessa Vernoud, Karine Gallardo-Guerrero

**Author notes:** These authors contributed equally (Vanessa Vernoud, Karine Gallardo-Guerrero). **Corresponding author**: Titouan Bonnot.

## Abstract

Sulfur availability affects crop yield, seed quality, and tolerance to environmental constraints. To understand how pea (*Pisum sativum*) leaves respond to sulfur deficiency alone or combined with moderate water deficit during the early reproductive phase, we employed a multi-omics approach. Sulfur deficiency reduced plant height, biomass and leaf carbon, and increased the nitrogen-to-sulfur ratio. Under this condition, 38 genes were up-regulated at both transcript and protein levels, including genes involved in sulfur metabolism and antioxidant responses, suggesting coordinated molecular adjustments that may mitigate low leaf sulfur status. Moderate water deficit alone had limited effects, but markedly altered plant growth, gene regulation and metal accumulation when combined with sulfur deficiency. Among synergistically up-regulated genes, twenty were linked to reactive oxygen species responses and activated early, while seven genes with sustained activation encoded glutathione *S*-transferases. This was associated with higher GST activity and likely contributed to limiting H_2_O_2_ accumulation in double-stressed leaves. One-third of differentially accumulated proteins were encoded by genes showing no transcriptional change under stress, including temperature-induced lipocalins with potential protective roles under combined stress. These findings enhance our understanding of multilevel molecular responses to stress interactions, which is essential for improving crop resilience under multi-stress conditions.

**Highlight:** Moderate water deficit amplifies molecular responses to sulfur deficiency in *Pisum sativum*, revealing synergistic responses at multiple layers of regulation under this stress combination.

## Introduction

Human activities are responsible for global changes such as global warming and climate change, some of which are now inevitable and irreversible (IPCC 2023). In addition to the increasing intensity and frequency of extreme events, the number and complexity of interacting factors affecting ecosystem functioning are rising rapidly (Paine et al. 1998; IPCC 2023). In the field, plants are often exposed to multiple growth-limiting stresses that can occur sequentially or simultaneously, such as elevated temperatures with water deficit (WD), or WD with nutrient deficiencies (Mittler 2006; Lecomte et al. 2023). Combinations of stresses can have more dramatic impacts on crop yields than individual stresses, as observed under combined heat and drought, or under drought and sulfur (S) deficiency (hereafter referred to as S-; Henriet et al. 2019; Cohen et al. 2021).

At the molecular level, studies performed under controlled or semi-controlled conditions have shown that combined stresses often induce non-additive effects and unique molecular responses, that cannot be predicted from single-stress responses (Rizhsky et al. 2002, 2004; Mittler 2006; Rasmussen et al. 2013; Zandalinas et al. 2021; Mahalingam et al. 2022; Tan et al. 2023). For example, ∼55% of genes responding to heat and/or drought stress in Arabidopsis displayed antagonistic or synergistic responses under combined heat and drought stress (Azodi et al. 2020). Synergy occurs when the effect of the stress combination exceeds the sum of individual effects. Such synergistic responses may reflect the activation of unique biological pathways that are critical for the plant tolerance to a given combination of stresses (Zandalinas et al. 2021), and must be considered for improving plant resilience to environmental constraints.

S is an essential macronutrient for plants. However, S- in soils is increasingly reported worldwide, due to factors such as stricter regulations on industrial SO_2_ emissions and the declining use of S-containing pesticides and S fertilizers (Haneklaus et al. 2008). This global trend results in S- symptoms in crops, affecting yields and seed quality (Scherer 2001; Haneklaus et al. 2008; Poisson et al. 2019; Borja Reis et al. 2021). In pea seeds and cereals such as wheat, the N:S balance strongly influences storage protein composition – a key determinant of seed quality (reviewed in Mondal et al. 2022; Bonnot et al. 2023). Besides its impact on the N:S ratio, S- affects the uptake and remobilization of other elements (Jacques et al. 2021). For instance, S- in Arabidopsis increases phosphate uptake, transport and accumulation in shoots (Allahham et al. 2020). Another well documented effect of S-on elemental accumulation is the increase in molybdenum contents (Shinmachi et al. 2010; Maillard et al. 2016; Jacques et al. 2021).

Sulfate is the major form of S available to plants. Under deficiency, the primary plant response includes the upregulation of several sulfate transporters (SULTRs) to enhance sulfate uptake and S availability for the metabolism. For example, the high-affinity transporters *SULTR1;1* and *SULTR1;2,* expressed in root epidermis and cortex, are significantly induced under S- in Arabidopsis (Takahashi et al. 2000; Vidmar et al. 2000; Yoshimoto et al. 2002; Maruyama-Nakashita et al. 2006). S- also triggers the accumulation of O-acetylserine (OAS) and induces a co-expressed set of genes known as the OAS cluster (Hubberten et al. 2012; Aarabi et al. 2016, 2021). Several members of the OAS cluster are also induced in wheat and pea seeds under S-limiting conditions (Bonnot et al. 2020; Henriet et al. 2021). Some studies suggest a role of this cluster in S sensing and signaling (Ristova and Kopriva 2022). Activation of molecular responses to S- in Arabidopsis is mediated by transcription factors (TFs) of the *ETHYLENE-INSENSITIVE3-LIKE (EIL)* family. The best characterized is SULFUR LIMITATION1 (SLIM1/EIL3), which functions as a key regulatory hub controlling the expression of multiple S deficiency-responsive genes including *SULTRs* and OAS-cluster genes (Maruyama-Nakashita et al. 2006; Apodiakou and Hoefgen 2023).

S also contributes to the production of antioxidant molecules like glutathione, and enzymes of the glutathione pathway such as glutathione-S-transferases and glutathione peroxidases. Under stress, reactive oxygen species (ROS) accumulate and can cause oxidative stress at high concentrations (see Mittler et al., 2022 for review). Thus, S- compromises ROS scavenging and redox homeostasis, altering the plant’s ability to tolerate additional stresses such as WD (reviewed in Samanta et al., 2020). Conversely, S supplementation can improve WD tolerance. Pre-treatment with appropriate level of S dioxide (SO_2_) increased survival and decreased hydrogen peroxide (H_2_O_2_) accumulation in WD-treated wheat seedlings (Li et al. 2021). Hydrogen sulfide (H_2_S) pre-treatment enhanced drought tolerance in Arabidopsis (Jurado-Flores et al. 2023), barley (Carrillo et al. 2025) and rice (Zhang et al. 2024).

We previously investigated the interaction between S- and WD during the early reproductive phase in pea (*Pisum sativum*; Henriet et al., 2019; Henriet et al., 2021). The combination of stresses had a synergistic negative impact on seed size and yield, yet WD mitigated the adverse effect of S- on seed storage protein composition. This suggests that under combined stress, pea plants may favor the production of fewer seeds with a well-balanced protein composition (Henriet et al. 2019, 2021). At the molecular level, fewer transcripts and proteins were differentially accumulated in developing seeds under combined stress compared with S- alone, indicating that key molecular responses activated under combined stress may occur in other tissues such as leaves, which are major nutrients sources for developing seeds.

To further dissect the molecular mechanisms activated under S- and WD, we performed a multi-omics analysis of pea leaves from the experiment described in Henriet et al. (2019, 2021). Transcriptomics, proteomics and ionomics data were obtained at different time points to identify molecular networks responding to each stress individually and in combination. Our results revealed a specific module of genes responding to S- (with or without WD) at both transcript and protein levels, involved in S metabolism and oxidative stress response. Combined S- and WD induced synergistic effects on the plant phenotype and on the transcriptome and proteome. Specific gene modules uniquely activated under combined stress were identified, involving genes playing a role in glutathione metabolism, or in metal ion or ROS responses. Our results also evidenced genes with protein-specific responses to stress, highlighting post-transcriptional and/or translational regulation under stress.

## Materials and methods

### Plant growth and treatments

Pea plants (*Pisum sativum* L., ‘Caméor’ genotype) were obtained from the experiment described in (Henriet et al. 2019). Briefly, germinated seeds were sown in pots filled with a mixture of perlite and sand (3/1, v/v). Plants were grown in a greenhouse, at 19°C/15°C (day/night) with a 16 h/8 h (light/dark) photoperiod. Plants were irrigated with a solution composed of: 4 mM KNO_3_, 2 mM Ca(NO_3_)_2_, 0.3 mM MgSO_4_, 0.9 mM MgCl_2_, 0.2 mM NaCl, 0.72 µM Na_2_MoO_4_, 0.10 mM EDTA-Fe-Na, 8.2 µM MnCl_2_, 1 µM CuCl_2_, 1 µM ZnCl_2_, 30 µM H_3_BO_3_,1 mM K_2_HPO_4_ (pH adjusted to 6.3 using H_3_PO_4_ before addition of K_2_HPO_4_). This solution provided 0.3 mM SO_4_^2-^ as the sole S source, a concentration previously established as sufficient for optimal growth (Henriet et al. 2019). After three weeks (5/6 node stage), S- treatment was applied to half of the plants: the substrate was rinsed twice using deionized water and twice using a S- nutritive solution depleted of MgSO_4_·7 H_2_O but containing 1.16 mM MgCl_2_, then plants were watered using this solution until the end of the experiment. After eight days following the beginning of S- treatment, all plants (including S-sufficient plants) at the 8-node stage (on the primary branch, secondary branches were removed across the experiment) were moved to an automated Plant Phenotyping Platform for Plant and Micro-organism Interactions (4PMI, Dijon, France). An excess of plants was initially grown for each condition (S-sufficiency and S-) to exclude those with highly heterogeneous development. Plants were weighted and watered four times a day to maintain a water-holding capacity of the substrate of 100%. At flowering of the second/third flowering nodes (*i.e.*, 16 days after the onset of S-), half of the plants (half of the S-sufficiency and half of the S deficiency-treated plants) were subjected to WD: irrigation was stopped to reach a water-holding capacity of the substrate of 50%, then this value was maintained for nine days. This level of irrigation corresponded to a leaf water potential of -1.3 MPa (Henriet et al., 2019). On the day of WD application, the first flowering node was tagged on all plants. Control and S-plants were at the same developmental stage at the time of WD imposition, as indicated by similar positions of the first flowering node and comparable total number of nodes (Supplementary Fig. S1). After nine days of WD, plants were rewatered normally, with their respective nutritive solution (S-sufficiency or S-). This experimental setup allowed to get four different sets of plants, grown under: control (no stress applied) condition, S-, WD, or a combination of S- and WD.

### Sampling and phenotypic measurements

A total of eight plants per condition and time point were grown in the 4PMI platform according to a randomized complete block design and used for phenotyping. Phenotypic measurements were performed on days 0, 2, 5, 9 and 12 after the start of the WD application. Note that day 12 corresponds to three days after the end of WD, *i.e.* three days of re-watering for the WD and WD/S conditions. For leaf area measurements, the leaves of each individual plant (*n* = 8) were spread out and scanned with an EPSON GT20000 (model J151A) scanner, and leaf area was measured using a custom image processing algorithm. Dry weight measurements were performed after drying tissues at 80°C for 48 h.

For -omics analyses, four biological replicates were arranged in the 4PMI platform according to a randomized complete block design. Each replicate consisted of leaves collected from the first two reproductive nodes of two plants grown side by side (*i.e*., a pool of leaves from two plants). Samples were collected in the morning, between 2h and 6h following the beginning of the light period, snap-frozen in liquid nitrogen and stored at -80°C. The same samples were used for transcriptomics, proteomics, ionomics and SCN content determination (see below).

### Osmotic potential estimation

Osmotic potential was estimated on the same plants as those used for the omics. For each biological replicate (*n* = 4), leaflets from the last fully expanded leaves of the two independent plants were collected, placed in a syringe, frozen in liquid nitrogen and stored at −80°C. After thawing, the cell sap was pressed out of a syringe, collected and centrifuged at 10,000 × g for 10 min at 4°C. The osmolality of the corresponding supernatant was measured using a vapour pressure osmometer (Wescor model 5520, Bioblock Scientific, Illkirch, France), and the osmotic potential (MPa) was then calculated according to the Van’t Hoff equation: π=(−R×T×osmolality of the extract), where T is the absolute temperature and R is the constant of perfect gas.

### Element analyses

S, C and N contents were determined from dried ground leaf samples (four biological replicates) using the Dumas method (Allen et al. 1974). Contents of C and N were determined from 5 mg of tissue powder on a Flash 2000 Elemental Analyzer (Thermo Fisher Scientific), and contents of S were determined from 20 mg of tissue powder mixed with 5 mg of tungsten trioxide on an elemental PYRO cube analyzer (Elementar). Two technical replicates per biological replicate were performed.

Elements referred to as ‘Ionomics data’ in the manuscript correspond to essential macro-nutrients (phosphorus, potassium, calcium, magnesium), micro-nutrients (iron, boron, manganese, zinc, copper, molybdenum, nickel) and other elements (cobalt, cadmium, lead, arsenic, vanadium), quantified by High Resolution Induced Coupled Plasma-Mass Spectrometry (HR ICP-MS, Thermo Scientific, Element 2TM). For each sample, 40 mg of dried ground leaf material were resuspended in 800 μL of concentrated HNO_3_, 200 μL H_2_O_2_ and 1 mL of Milli-Q water. All samples were then spiked with three internal standard solutions containing gallium, rhodium and iridium with final concentrations of 5, 1 and 1 μg L^−1^, respectively. After microwave acidic digestion (Multiwave ECO, Anton Paar, les Ulis, France), all samples were diluted with 50 mL of Milli-Q water to obtain solutions containing 2.0% (v/v) nitric acid. Before HR ICP-MS analysis, samples were filtered through a 0.45 μm teflon filtration system (Digifilter, SCP Science, Courtaboeuf, France). Quantification of each element was performed using external standard calibration curves and concentrations were expressed in μg.g^-1^ of dry weight. Data are presented in Supplementary Dataset S1.

### Quantification of H_2_O_2_ levels

Hydrogen peroxide (H_2_O_2_) levels in leaves were measured by acid extraction as described by Noctor et al. (2016) with few modifications. Briefly, 100 mg of nitrogen-ground pea leaves were homogenized with 1 mL of 0.2 M HCl and thawed on ice before centrifugation (16,000 × g, 10 min, 4°C). 500 µL of each supernatant were mixed with 100 µL of 0.2□M phosphate buffer (pH□5.6) and adjusted at pH 5 using 0.2 M NaOH. To remove ascorbate, 300 µL of this mixture were incubated for 30 min with 2.4 units (6 µL) of ascorbate oxidase (Merck-Sigma-Aldrich). In a 96 well plate, 200 µL of 500 µM luminol working solution, prepared by diluting a 150 mM stock solution (in DMSO) into 0.2 NH3 buffer (pH 9.5), was mixed with 25 µL of ascorbate oxidase-treated sample supernatant. In parallel, a standard curve was prepared containing 0 to 5□nmol H_2_O_2_ in 200 µL of 0.2 M NH3 (pH 9.5) for each sample. Finally, 40 µL of 500 µM K3Fe(CN)6 (Merck-Sigma-Aldrich) were injected to each samples using the TECAN Infinite 200 Pro multimode plate reader, mixed rapidly, and light emission was recorded over 2 s. H_2_O_2_ production was expressed in nmol of H_2_O_2_ per mg of fresh weight. Assays were performed in three technical replicates with three independent biological experiments.

### Measurement of GST activity

The activity of glutathione S-transferase (GST) was determined according to Czékus et al. (2020) with some modifications. Briefly, 250 mg of nitrogen-ground pea leaves, supplemented with 10 mg of polyvinylpolypyrrolidone (PVPP), were homogenized in 1 mL of cold extraction buffer (50 mM sodium-phosphate pH 7.0, 1 mM phenylmethylsulfonyl fluoride, 1 mM EDTA). The mixture was thawed on ice with regular vortexing. Homogenates were then centrifuged (25,000 × g, 30 min, 4°C) and the resulting supernatants were used directly for GST activity assays. In a 96-well plate, 180 µL of extraction buffer containing 100 mM reduced glutathione (GSH) was added to 10 µL of supernatant containing GST. The initial absorbance at 340 nm (OD₃₄₀) was measured using a TECAN Infinite 200 Pro at 25°C. The enzymatic reaction was initiated by adding 10 µL of 20 mM 1-chloro-2,4-dinitrobenzene (CDNB) to each sample, and OD₃₄₀ was recorded every 20 seconds for 5 minutes. In parallel, protein concentration in each sample was determined using the Bradford method. The enzyme reaction rate was calculated from the linear portion of the OD₃₄₀ curve over time, using the CDNB extinction coefficient (0.0096 µM⁻¹cm⁻¹) and the pathlength of the solution in the well (0.597 cm). Activity was expressed as the number of nmoles of GSH conjugated to CDNB per minute per mg of total protein. The assay was performed in three independent experiments.

### Shotgun proteomics

Protein sample preparation, analysis by LC-MS/MS and quantification, were performed as described in (Henriet et al. 2021). Briefly, total proteins were extracted from 110 mg of ground leaf tissues. Protein samples were lyophilized, then solubilized in buffer containing 6 M urea, 2 M thiourea, 10 mM DTT, 30 mM Tris-HCl pH 8.8 and 0.1% zwitterionic acid labile surfactant. For each sample, 20 µg of proteins were alkyled then digested overnight at 37°C using trypsin. Trypsin-digested proteins were desalted and analyzed by LC-MS/ MS using an Eksigent nlc425 device coupled to a Q Exactive mass spectrometer (ThermoFisher Scientific), with a glass needle (non-coated capillary silica tips, 360/20-10, New Objective). Proteins were identified using X!Tandem v.2015.04.01.1 (Craig and Beavis 2004), by matching peptides against the *P. sativum* v.1a database (https://urgi.versailles.inra.fr/jbrowse/gmod_jbrowse/; (Kreplak et al. 2019). Identified proteins were filtered and grouped using the X!TandemPipeline software v.3.4.3 (Langella et al. 2017). Relative protein quantification was performed using the MassChroQ software v.2.2 (Valot et al. 2011). Protein quantification data is provided in Supplementary Dataset S3.

### RNA extraction and sequencing

Extraction of RNAs from leaf tissues, RNA-sequencing (RNA-Seq) and data processing were performed as described in (Henriet et al. 2019). Briefly, total RNAs were extracted from ground leaf tissues using an RNeasy Plant Mini Kit (QIAGEN), then a DNAse treatment with an RNase-Free DNase Set (QIAGEN) and a purification step using lithium chloride precipitation were performed. RNA quality was checked on a 2100 Bioanalyzer (Agilent Genomics). RNA-Seq libraries were prepared using an Illumina TruSeq Stranded mRNA sample prep kit, and 11 PCR cycles were performed to amplify the libraries. Library quality was checked using a Fragment Analyzer (Agilent Genomics) and libraries were sequenced on the Illumina HiSeq3000 to obtain 2×150 bp paired-end reads. Quality of raw reads was assessed using the FastQC v0.11.2 software (https://www.bioinformatics.babraham.ac.uk/projects/fastqc/). Trimming of low-quality and adapter sequences was performed on raw reads using Trimmomatic v0.32 (Bolger et al. 2014). Trimmed reads with less than 25 bp and unpaired reads were removed. Filtered reads were mapped to the *P. sativum* v1a reference genome (https://urgi.versailles.inra.fr/Species/Pisum/Pea-Genome-project), using the alignment program HISAT2 v2.0.5 (Kim et al. 2015). Read counting was performed using FeatureCounts v1.5.0-p3 (Liao et al. 2014) on gene annotation v1b (Henriet et al. 2019). Only genes with at least 10 reads total in the experiment were considered as expressed and were used for downstream analyses (29575 genes in total). Gene expression was normalized using the VST transformation proposed by the DESeq2 package (Love et al. 2014). Gene expression data is provided in Supplementary Dataset S3.

### RT-qPCR

Five µg of purified RNA were reverse-transcribed with the iScript cDNA synthesis kit (Bio-Rad) according to manufacturer’s protocol in a final volume of 20 µL. Real-time PCR was performed on a LightCycler 480 (Roche) using the GoTaq qPCR Master Mix (Promega, Madison, USA) with 3 µL of 50 times diluted cDNA, 0.2 µL of each primer (10 µM) in a final volume of 10 µL, and the following program: 95°C for 5 min; and 40 cycles of 95°C for 10 s and 60°C for 1 min. A melting curve analysis was performed by a further cycle from 50°C with a 0.11°C increase per second up to 95°C, and the fluorescence was measured every 0.5°C. Reactions were performed in duplicates from each biological replicate. Expression levels were calculated using the ΔΔCT method (Schmittgen and Livak 2008), with α-Tubulin (*Psat5g091400*) chosen as the reference gene based on RNA-Seq data indicating stable expression across our samples. The specificity of the primer pairs was checked by melting curve analysis. Primer pair efficiency was verified with a 5-point dilution series of Pea Caméor genomic DNA. Primer sequences are listed in Supplementary Fig. S2.

### Statistical analyses

#### Identification of variables significantly affected by treatments

All statistical analyses were performed with the software program R v. 4.1.2 (R Core Team 2022). To assess the influence of S- at the beginning of the experiment on phenotypic variables, on leaf CNS contents, and on the leaf osmotic potential, unpaired Student’s t-tests were performed to compare treatments S- and control at day 0. Significant difference between treatments was judged at *P*-value < 0.05.

For data collected from days 2 to 9, the effects of time, treatments and the interaction between these main effects were analyzed using the following model: design = ∼ Time + Treatment + Time:Treatment. For transcriptomics, differential expression analysis was performed from raw counts of the 29575 filtered genes that were considered as expressed (see above). Likelihood ratio tests (LRTs) were performed with the DESeq2 package (Love et al. 2014), using the model described above. The LRT is conceptually similar to an analysis of variance (ANOVA) calculation in linear regression (Love et al. 2014). For other -omics data, two-way ANOVAs were performed for each individual variable using the aov() function. For all data, significance of factors was judged at *P* < 0.05 after FDR correction using the Benjamini–Hochberg procedure (Benjamini and Hochberg 1995). Comparisons of treatments were then performed at each time point. For transcriptomics, pairwise comparisons were realized using the DESeq2 package (Love et al. 2014). For other omics data, Tukey’s honestly significant difference (HSD) mean- separation tests were performed, using the TukeyHSD() function. Variables were considered as differentially responding to treatments at a given time point (*e.g.* in response to S- as compared to control at day 2) if they showed *i*) a significant treatment effect (FDR < 0.05, LRTs or ANOVAs) and *ii*) a significant difference between treatments (FDR < 0.05, pairwise comparisons for transcriptomics; adjusted P-value < 0.05, Tukey’s HSD tests for other omics data). To consider as significant both variables with high or low magnitude of change in response to stress, no cutoff was applied on Fold Change values.

For data obtained at day 12, the same procedure was employed as for the analysis at days 2-9, with the following modification in the model used for LRTs (transcriptomics) and ANOVAs (other -omics data): design = ∼ Treatment. Statistical results are provided in Supplementary Dataset S2.

#### Enrichment analyses

Gene Ontology (GO) enrichment analyses were conducted using the ‘enricher’ function of clusterProfiler R package v. 4.14.6 (Yu et al. 2012). For GO enrichment in the M16-MP1 overlapping genes, the gene universe corresponded to GO terms associated with overlapping genes between the mRNA and protein network. For GO enrichment in modules M2, M7 and M9, the gene universe corresponded to GO terms associated with all genes from the mRNA network. GO terms with an adjusted *P*-value < 0.05 were considered as significantly enriched in the selected subset of genes.

To identify the enrichment of specific sets of genes (with a time, treatment or time×treatment effects, or with a synergistic response under the combined stress condition) within network modules, Fisher’s exact tests were performed. Proportions of these specific sets of genes within each module were compared to their proportions in all genes found in the network. Significant differences were judged at *P* < 0.05.

#### Correlation between proteomics and transcriptomics data

For each expressed gene for which a protein was successfully quantified in shotgun proteomics (2240 genes in total), correlations were calculated between protein and mRNA data, with the R cor() function and the Spearman method. Correlation coefficients are provided in Supplementary Dataset S4.

### Co-expression network analysis

Weighted Gene Co-expression Network Analyses were conducted with the R package ‘WGCNA’ v. 1.72-5 (Langfelder and Horvath 2008). Two separate analyses were performed, from data obtained at the 1) transcriptome and 2) proteome levels. Prior to analysis, transcriptomic data were filtered to remove genes with low expression values that could introduce noise into the network analysis. Genes with raw counts > 10 in at least 50% of the samples were considered for the analysis, which represents 21281 genes. Normalized genes expression values (after VST transformation) were then used. No filtering was applied on proteomic data, all 2261 quantified proteins were used for the analysis. Data obtained at days 0 (control and S- conditions), 2, 5, 9 and 12 were used. Adjacency matrices were built using a soft threshold power of 12 and 16 for the analyses performed from transcriptomic and proteomic data, respectively. To identify modules of co-expression, the minimum module size was set at 30. Similar modules were merged, using a dissimilarity threshold of 0.25. Module eigengene values were used to represent module expression patterns. Module information is provided in Supplementary Dataset S7. To identify pea TFs, the iTAK program was run on proteins of the v1b pea genome annotation (Zheng et al. 2016). Results are provided in Supplementary Dataset S8. For selected pea genes, homologous genes in Arabidopsis were identified using reciprocal BLASTP search (https://plants.ensembl.org/Multi/Tools/Blast). The phylogeny of sulfate transporters (SULTRs) was built from protein sequences, using the interactive phylogenetics module of the Dicots PLAZA v5 database (https://bioinformatics.psb.ugent.be/plaza/versions/plaza_v5_dicots/). The software MUSCLE v. 3.8.31 and FastTree v. 2.1.7 were used for the multiple sequence alignment and the phylogenetic tree, respectively.

### Data visualization

Heatmaps were generated with the R package ‘pheatmap’ v. 1.0.12 (Kolde 2019). Venn diagrams were drawn using the R package ‘eulerr’ v. 7.0.1 (Larsson 2022). Enriched GO terms were represented as described in (Bonnot et al. 2019). Networks were visualized with the software ‘CYTOSCAPE’ v. 3.9.0 (Smoot et al. 2011), using a Prefuse Force Directed Layout. Links with a weight > 0.15 (mRNA network) or a weight > 0.10 (proteome network) were used for visualization. All other plots were prepared using the R package ‘ggplot2’ v. 3.4.4 (Wickham 2016).

## Results

### Water deficit combined with S deficiency severely affects the plant growth

To explore the interplay between WD and S-, pea plants were exposed to S- starting three weeks after sowing (5/6 leaf stage), to moderate WD (50% of the substrate water holding capacity) applied at the onset of flowering for nine days, or to the combination of both WD and S- (WD/S-, Fig. 1A). S- was imposed 16 days before WD to ensure a reduced leaf S status at the time of WD imposition. The experimental setup is detailed in Henriet et al. (2019, 2021). Phenotypic variables were measured across plant compartments, and leaves from the first two reproductive nodes were harvested from control and stressed plants at days 0 (for control and S- conditions only), 2, 5, 9 and 12 to conduct -omics analyses (Fig. 1). Raw data and statistical results are presented in Supplementary Datasets S1 and S2, respectively. For WD and WD/S- conditions, day 12 corresponded to three days after rewatering, and informs on the plant recovery after WD. Separate statistical analyses were performed for data obtained at day 0 and day 12 due to different treatment modalities compared to other time points (see Methods).

**Fig. 1.**
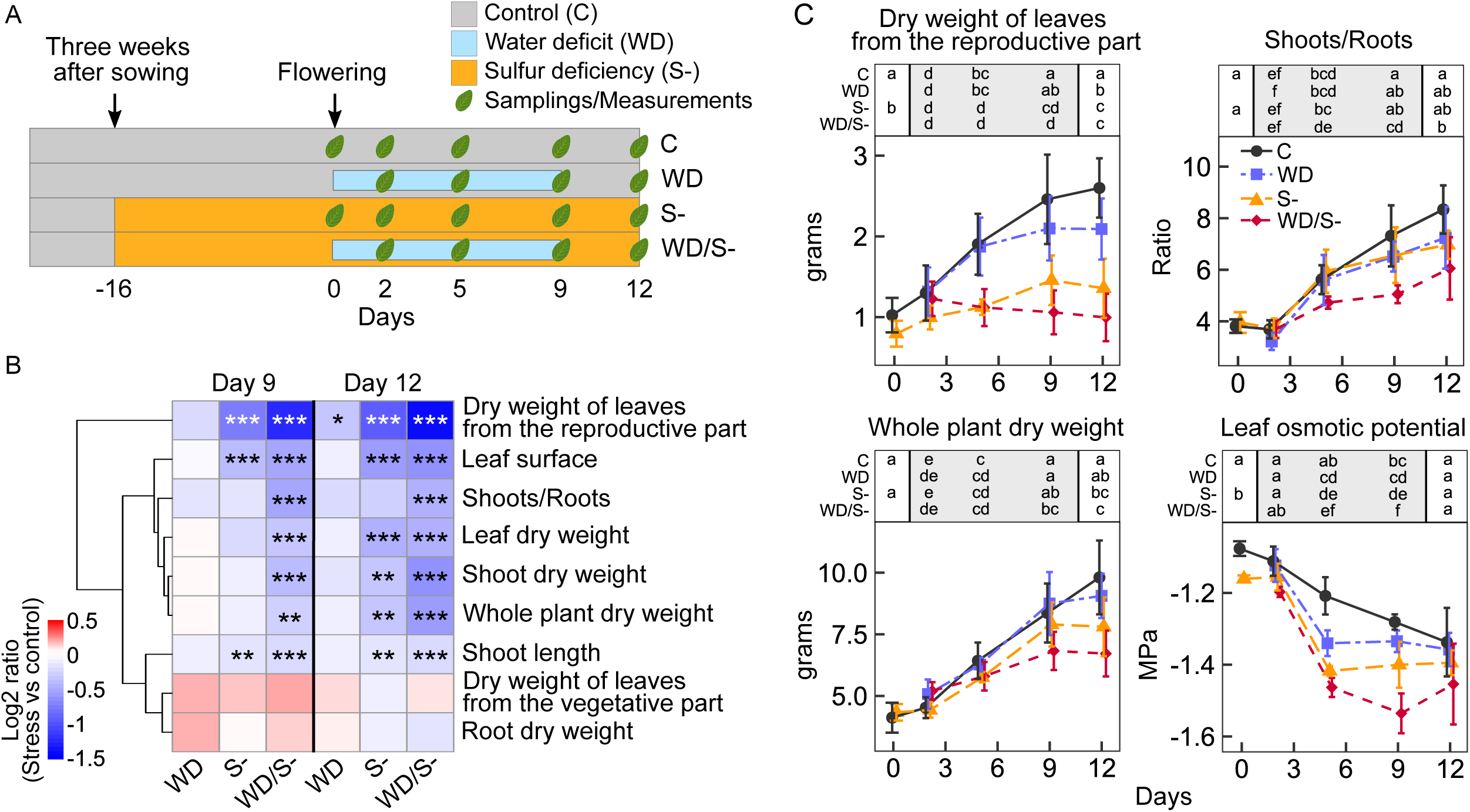
Influence of water deficit (WD), sulfur deficiency (S-) and WD/S- on pea plant phenotypic variables. A, Experimental design. Days 0 and 9 correspond to the beginning and end of WD, respectively. For the conditions WD and WD/S-, day 12 therefore corresponds to three days of rewatering for plant recovery. B, Effects of stresses on phenotypic variables, at day 9 and day 12. Data are represented as heatmaps, blue and red representing lower and higher values in the stress condition as compared to control, respectively. Variables with similar responses to stress are grouped using a hierarchical clustering method. Data are Log2 ratio (stress condition vs control) calculated from means of n = 8 biological replicates (eight individual plants per condition). In B, asterisks indicate significant differences between stress and control conditions (*, P < 0.05; **, P < 0.01; ***, P < 0.001). C, Profiles of selected phenotypic variables and of osmotic potential in leaves. Data are means ± S.D. of n = 8 and n = 4 biological replicates for phenotypic variables and for osmotic potential, respectively. In C, above each plot, different letters represent significant differences between treatments. Three separate statistical analyses were performed: comparison of S- and control at day 0 (Student’s t-test); comparisons of all treatments from day 2 to day 9 (two-way ANOVA followed by Tukey’s HSD tests); comparisons of all treatments at day 12 during the plant recovery (one-way ANOVA followed by Tukey’s HSD tests, see methods for details). Raw data and statistical results are presented in Supplementary Datasets S1 and S2, respectively.

WD applied alone had only minor effects on the measured traits (Fig. 1B). The only significant changes were a decrease in the dry weight of leaves from the reproductive part of the plant at day 12 (Fig. 1, B and C), and a lower osmotic potential at day 5 (-1.34 MPa under WD, -1.21 MPa under control; Fig.1C).

Under S-, several traits were significantly affected, including reduction in leaf surface area, shoot length, and leaf, shoot and whole-plant dry weight (Fig. 1B and 1C; Supplementary Fig. S3). Together with a significant reduction in the leaf osmotic potential (days 0, 5 and 9), these results highlight the negative impact of S- on the plant development and physiology (Fig. 1C). Of note, leaf osmotic potential was significantly reduced at day 0 under S-, before WD imposition.

When both stresses were combined (WD/S-), several traits were specifically or more severely affected than under single stress conditions (Fig. 1B and 1C). The shoot-to-root ratio decreased specifically under WD/S- at days 9 and 12, whereas no significant effect was observed under single stress conditions (Fig. 1C). This was associated with a significant decrease in the shoot dry weight (Supplementary Fig. S3). Whole-plant dry weight was significantly reduced compared to control at both days 9 and 12 (by 31.5% at day 12; *P* ≈ 6.14E-05), whereas under single stress, a significant effect was only observed under S- at day 12 (Fig. 1C). A greater reduction in leaf osmotic potential was also observed under WD/S- at day 9 (Fig. 1C). Taken together, these results showed that while WD alone had moderate effects, it severely impaired plant growth when combined with S-.

### Single and combined stress treatments modify the leaf elemental composition

To estimate the impact of single or combined stress conditions on the plant nutritional status, we next quantified C, N and S contents in leaves using the Dumas method. No significant differences were observed in response to WD alone, although leaf N content tends to be reduced under this condition (Fig. 2A). Both S- and WD/S-significantly reduced leaf C, N and S contents at one or more time points (Fig. 2A). At day 0, all three elements showed reduced accumulation in response to S-, indicating that leaf elemental composition was altered before WD imposition, as expected from our experimental design where S- was applied at the 5/6 leaf stage, *i.e.* about 16 days before the onset of WD. The reduced C accumulation likely reflects impaired photosynthetic capacity under S-, as S is required for photosynthetic components including iron-sulfur cluster-containing proteins of the electron transport chain. Leaf S content was remarkably reduced under S- and WD/S-, with no clear distinction between control and WD, or between S- and WD/S-, highlighting that our S-treatment strongly reduced S accumulation and/or translocation in pea leaves, regardless of water supply. Balance between C, N and S contents were also altered by treatments, especially the N/S ratio, which highly increased under both S- and WD/S- (Fig. 2A).

**Fig. 2.**
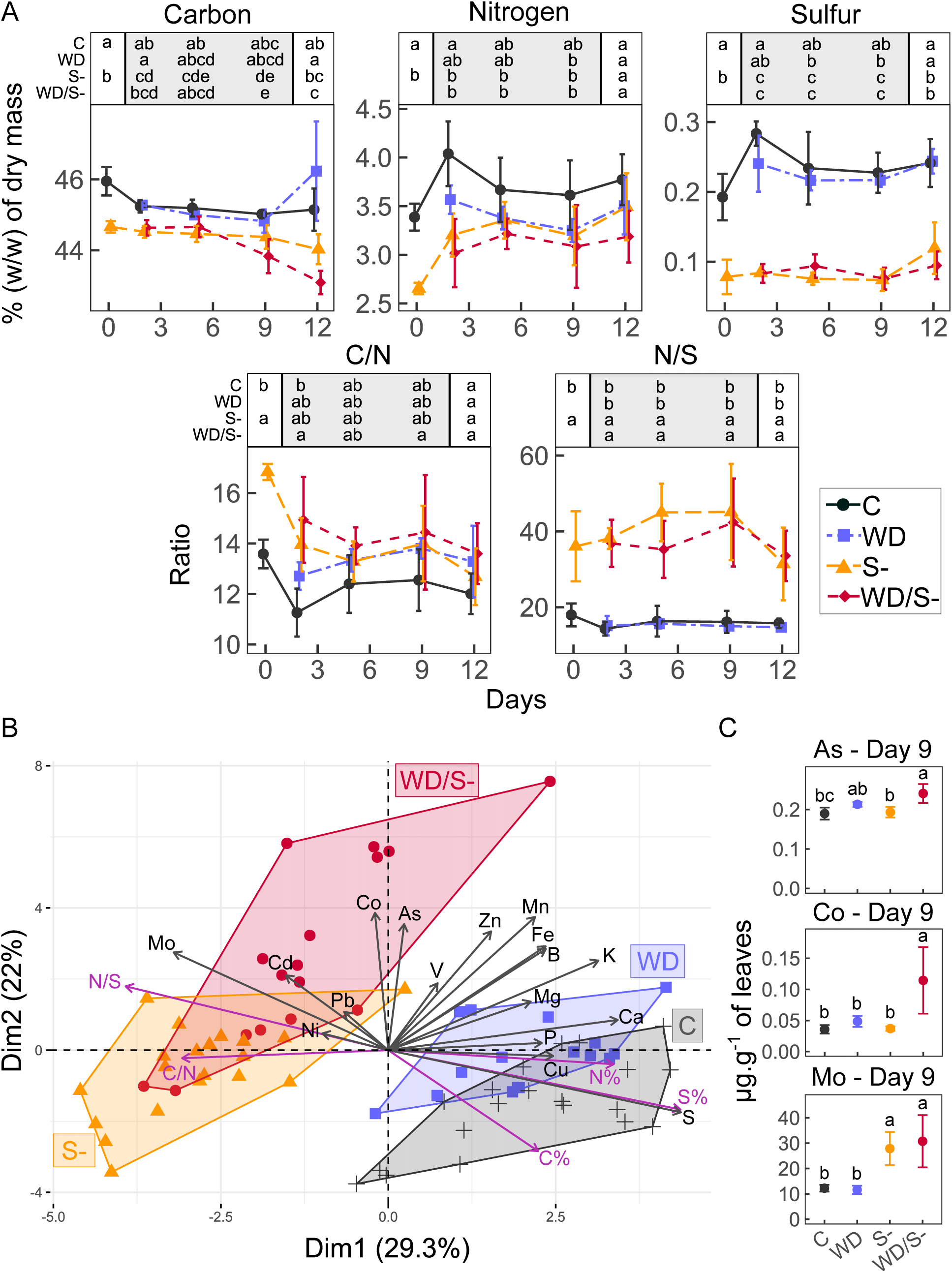
Elemental composition of pea leaves in response to water deficit and/or sulfur deficiency. A, Profiles of accumulation of Carbon (C), Nitrogen (N), Sulfur (S), as well as C/N and N/S ratios (means ± S.D, n = 4 biological replicates). Above each plot, different letters represent significant differences between treatments. Three separate statistical analyses were performed: comparison of S- and control at day 0 (Student’s t-test); comparisons of all treatments from day 2 to day 9 (two-way ANOVA followed by Tukey’s HSD tests); comparisons of all treatments at day 12 during the plant recovery (one-way ANOVA followed by Tukey’s HSD tests, see methods for details). B, Principal component analysis of elements presented in A (purple arrows) and in Supplementary figure S4 (black arrows). The convex hulls group samples by treatment. C, Profiles of selected elements at day 9 (means ± S.D, n = 4 biological replicates). Full profiles of all quantified elements are presented in Supplementary Fig. S4).

To further evaluate the effects of treatments on the plant nutritional status, we extended our analyses to other essential macro-nutrients (phosphorus, potassium, calcium, magnesium), micro-nutrients (iron, boron, manganese, zinc, copper, molybdenum, nickel) and elements (cobalt, cadmium, lead, arsenic, vanadium) using high-resolution inductively coupled plasma mass spectrometry (HR ICP-MS, Fig. 2B; Supplementary Fig. S4). PCA analysis from this dataset combined with C, N and S data evidenced two distinct groups of samples on the first dimension: C and WD opposed to S- and WD/S- treatments (Fig. 2B). This indicates a strong influence of S-on the leaf elemental composition. Notably, significant reduction of calcium, iron, potassium, magnesium and phosphorus were observed under S- compared to control (Supplementary Fig. S4). Molybdenum showed remarkable higher contents in leaves under S- and WD/S-, compared to control and WD conditions (Fig. 2C; Supplementary Fig. S4).

Dimension 2 of the PCA separates WD/S- from other treatments, suggesting distinct patterns of element accumulation under stress combination. Significantly, iron and zinc showed higher accumulation under WD/S- compared to control or to single stress conditions, at day 9 and during rewatering at day 12 (Supplementary Fig. S4). Similarly, cobalt showed higher accumulation at day 9 under WD/S-. The non-essential element arsenic, was also more accumulated in response to WD/S- (Fig. 2C; Supplementary Fig. S4). Increased accumulation of cadmium in leaves was specifically detected in response to S- compared to control at day 2 (Supplementary Fig. S4). While the accumulation of non-essential elements such as arsenic and cadmium was detected in leaf tissues, it is important to note that these elements were not intentionally supplied and likely represent trace contaminants originating from the growth substrate (perlite/sand mixture), as the nutrient solutions were prepared with deionized water.

Together, these results showed that stress treatments, especially S- and WD/S-, led to perturbations of the leaf elemental composition. Molybdenum and N/S ratio, which both showed strong negative correlation with leaf S content (Fig. 2B), were therefore indicators of S deprivation. Arsenic and cobalt, which particularly accumulated under WD/S-, seemed to be more specific to combined stress.

### Treatment-driven modules of transcripts and proteins

To identify genes and biological pathways differentially regulated under WD and/or S-, we analyzed the leaf transcriptome and proteome using mRNA-Sequencing and shotgun proteomics, respectively. In total, 29572 genes were expressed and 2261 proteins were quantified (Fig. 3A; Supplementary Dataset S3). Since day 12 corresponded to the recovery period after WD, omics data obtained at this time point were statistically analyzed separately (Fig. 3A).

**Fig. 3.**
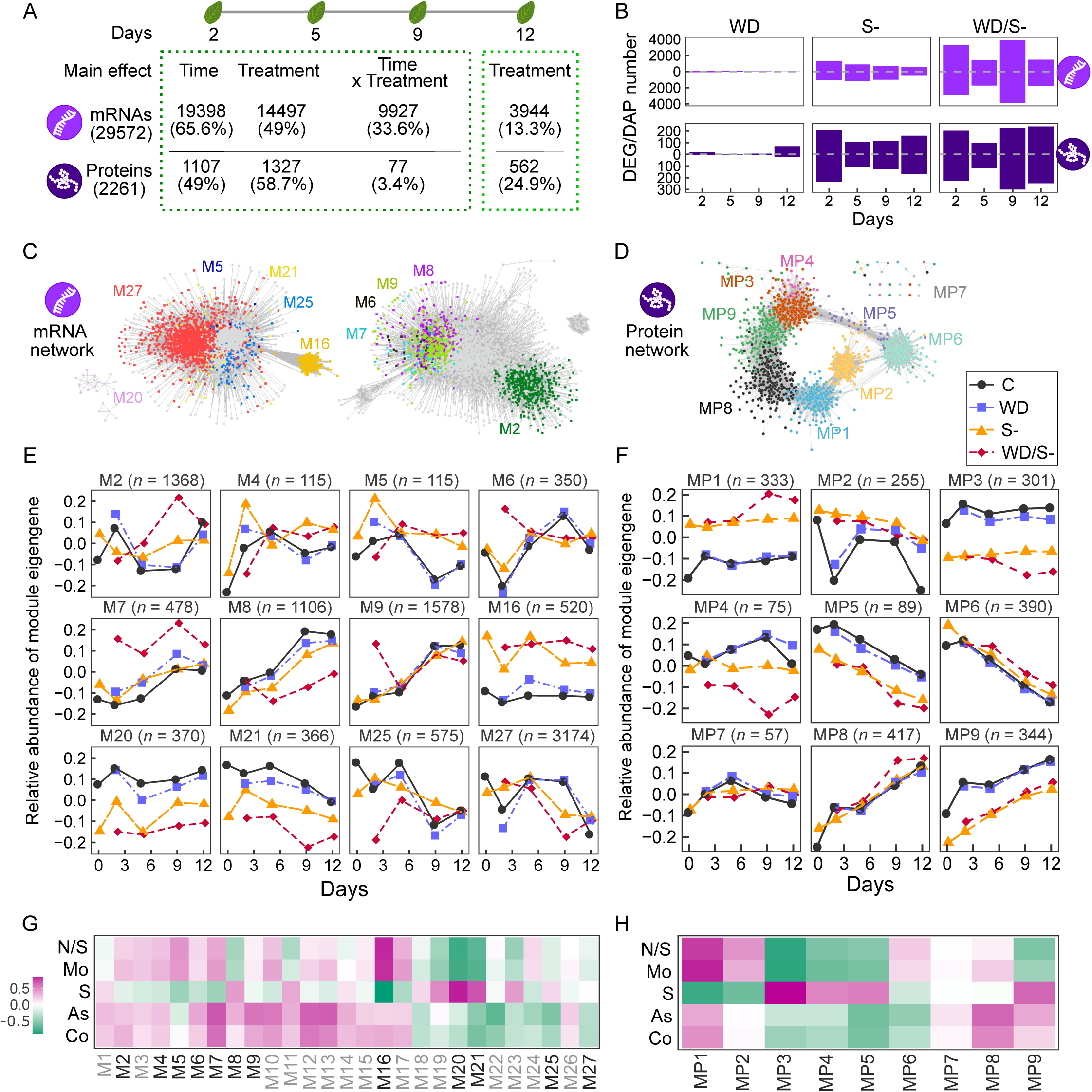
Transcript and protein modules of response to water deficit and/or sulfur deficiency in pea leaves. A, Results of statistical analyses showing the number of mRNAs and proteins with a significant (FDR < 0.05) effect of time, treatment, and the interaction between time and treatment (Time × Treatment). Two separate analyses were performed, one for day 2 to day 9, for which conditions were the same (Two-way ANOVA), and one for day 12, during the plant recovery following WD (One-way ANOVA). The percentage of analyzed molecules with a significant effect is indicated in parentheses (e.g. the accumulation of 49% of the proteome is significantly affected by time). B, Barplots showing the numbers of differentially accumulated mRNAs and proteins in response to the stress conditions as compared to control, at each developmental stage. This results from pairwise comparisons following the analysis presented in A (variables with a significant treatment effect were used for pairwise comparisons). C and D, Co-expression networks generated with WGCNA and visualized in Cytoscape using a Prefuse Force Directed layout, with an edge threshold cutoff of weight > 0.15 (mRNA network, C) and 0.1 (Protein network, D). E and F, Profiles of module eigengenes from networks presented in C and D, respectively (means ± S.D, n = 4 replicates). For the mRNA network, only modules enriched for genes with a significant treatment effect are colored in C and represented in E. Profiles of all module eigengenes are presented in Supplementary Fig. S5 and results of the enrichment analysis are presented in Supplementary Fig. S6. Note that module M4 is not represented in C, because of the edge threshold cutoff used for the network visualization. E and H, Heatmaps representing correlations (Spearman) between trait data (elements and osmotic potential) and module eigengenes of mRNA (E) and protein (H) networks. Green and pink colors indicate negative and positive correlations, respectively.

Analyses of variance at days 2-9 revealed that treatments significantly (FDR < 0.05) altered the expression of 14497 (49%) genes and the accumulation of 1327 (58.7%) proteins (Fig. 3A). Additionally, 33.6% of the transcriptome showed a significant time × treatment interaction effect, meaning that responses to treatments are influenced by time for these genes (Figure 3A). RT-qPCR was performed for five selected genes that showed a significant time, treatment and/or time × treatment effect (*Psat4g082000*, *Psat3g037400*, *Psat5g207000*, *Psat1g179680*, *Psat3g042160*). A strong correlation (*R* = 0.88, *P* < 2.2e-16) was observed between the RNA-Seq and RT-qPCR data, thus validating our RNA-Seq data analysis (Supplementary Fig. S2). The proportion of proteins with a significant time × treatment interaction effect was much lower (3.4%), suggesting smaller variations in the stress response over time for proteins.

Next, pairwise comparisons performed on variables with a significant treatment effect (FDR < 0.05) revealed that many more mRNAs and proteins were differentially accumulated in response to S- and WD/S- than to WD alone (Fig. 3B). We observed that numbers of differentially expressed genes (DEGs) and, to a lesser extent, of differentially accumulated proteins, were larger under WD/S- than under single stresses (Fig. 3B). For instance, at day 9, 3901 up-regulated DEGs were identified under WD/S-, whereas only 23 and 709 genes were up-regulated in response to WD and S-, respectively. The number of differentially accumulated molecules also differed depending on the days. For instance, under WD/S-, 3286, 1430 and 3901 genes and 202, 98 and 225 proteins were up-regulated at days 2, 5 and 9, respectively (Fig. 3B).

To further capture the influence of treatments on gene expression, we employed a weighted-gene co-expression network approach, from data obtained at the transcriptome level (mRNA network) and at the proteome level (protein network). Resulting mRNA and protein networks were composed of 27 and nine modules, respectively, exhibiting different patterns over time and/or in response to treatments (Fig. 3C-3F; Supplementary Fig. S5). In the mRNA network, we focused on 12 modules that were enriched for genes with a significant treatment effect (identified in Fig. 3A and Supplementary Fig. S6), and that we defined as treatment-driven modules (Fig. 3E).

Correlations between modules and selected elements (cobalt, arsenic, S, molybdenum and N/S ratio) were calculated. The mRNA M16 and protein MP1 modules were strongly negatively correlated with S content, while positively correlated with molybdenum and N/S (Fig. 3G-3H). Both modules showed high transcript (M16) and protein (MP1) accumulation under S- and WD/S-, relative to control and WD, across all time points (Fig. 3E-3F). In contrast, modules M20 and MP3 displayed the opposite trend, being down-regulated under S- and WD/S-. Together, these results suggest that modules M16, M20, MP1 and MP3 may be primarily driven by S availability.

Modules M7 and MP8 were positively correlated with arsenic and cobalt, and contained mRNAs and proteins that accumulated more strongly under WD/S-. Interestingly, the mRNA network revealed additional modules with responses specific to WD/S- at particular time points, such as M2 (day 9) and M9 (day 2, Fig. 3E). Such time-specific responses were less evident in the protein network.

These results provided the basis for the following sections, where we investigated the processes and genes that may mitigate the effects of S- with and without WD.

### A group of genes activated by S deficiency at both mRNA and protein levels

Our network analysis identified modules M16 (mRNAs) and MP1 (proteins) as consistently activated under S- and WD/S- treatments. Two SULTRs (*Psat2g074400* and *Psat3g185920*) were assigned to M16. *Psat2g074400* is homologous to AtSULTR of group 1 involved in sulfate uptake and source to sink transport in Arabidopsis (Yoshimoto et al. 2002) and *Psat3g185920* corresponds to *PsSULTR4* involved in vacuolar sulfate efflux (Bachelet et al. 2024; Supplementary Fig. S8).

Since M16 and MP1 showed similar patterns, we asked to what extent transcriptome responses were conserved at the protein level. To address this, we compared mRNA and protein profiles for the 2240 genes analyzed in both datasets. Across all treatments and time points, correlations between transcript and protein abundance ranged from -0.67 to 0.92, with a median of 0.12 (Fig. 4A). This overall low correlation suggests that post-transcriptional regulation strongly influences mRNA translation for most genes. Nonetheless, we identified a subset of 308 genes (13.7%) with correlations >0.5, that may be primarily regulated at the transcriptional level. Among all modules, M16 exhibited the highest overall transcript-protein correlation (median = 0.672 vs 0.119 for other modules; Fig. 4B; Supplementary Fig. S7).

**Fig. 4.**
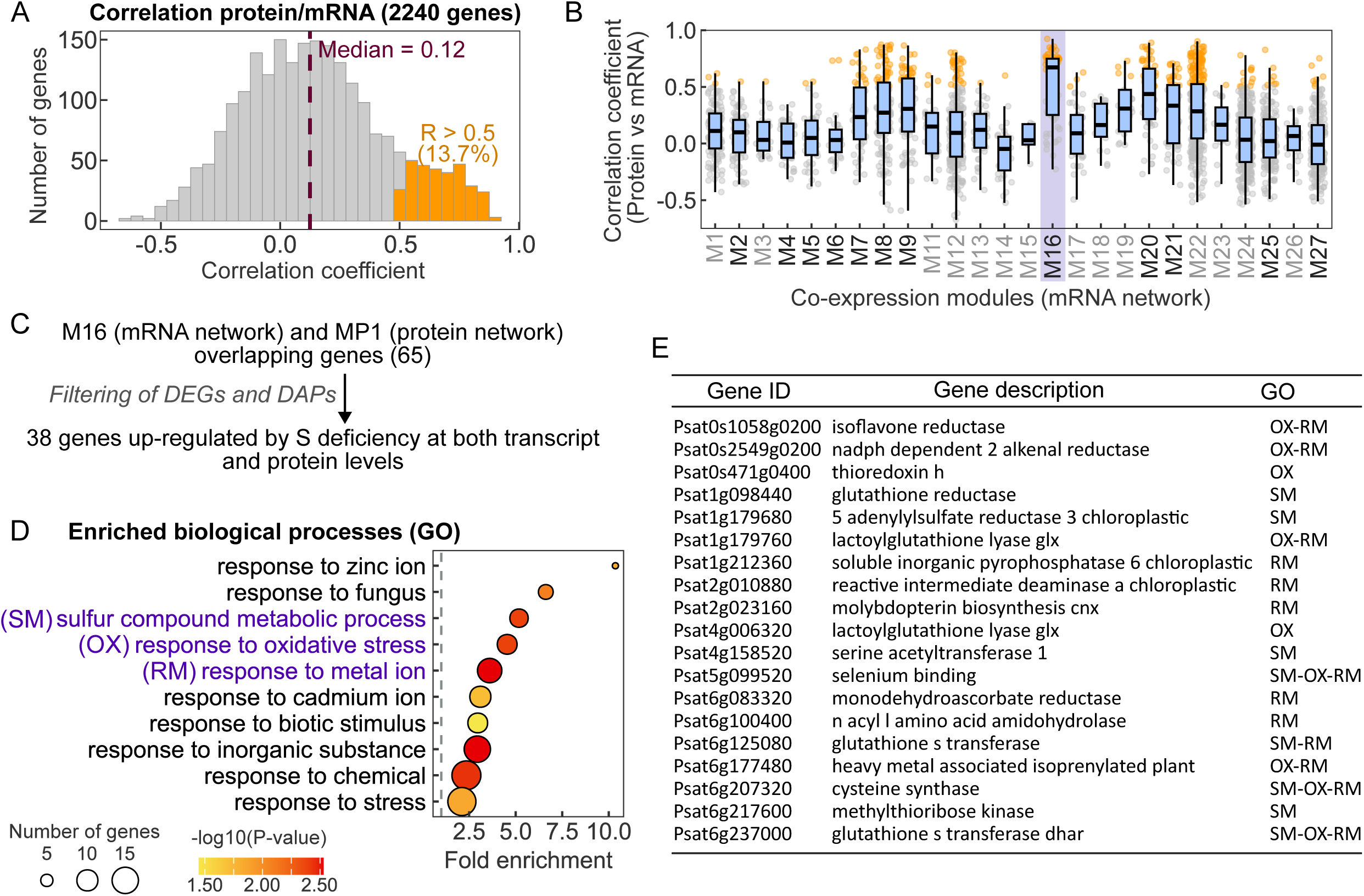
Identification of up-regulated genes under sulfur deficiency at both transcript and protein levels in pea leaves. A, Histogram showing the distribution of correlation (Spearman) coefficients for the comparisons between proteins and mRNAs (data obtained at days 2, 5, 9 and 12 from all treatment conditions). Genes with a R > 0.5 (protein vs mRNA) are highlighted in orange. B, Repartition within modules of genes analyzed at both transcript and protein levels (presented in A), with boxplots showing the distribution of correlation (Spearman) coefficients for the comparison protein vs mRNA. Dots represent individual genes. Genes with a R > 0.5 (Protein vs mRNA) are highlighted in orange. Boundaries of the boxes represent the 25th and 75th percentiles, and horizontal lines within boxes represent medians. Note that no data are represented for module M10, indicating that no corresponding protein was quantified for any of the genes in this module. C, Selection of genes up-regulated by sulfur deficiency at both transcript and protein levels. D, Top 10 enriched (lowest P-values) biological processes in the list of 38 M16 DEGs and MP1 DAPs overlapping genes. E, List of genes with a Gene Ontology annotation corresponding to ‘sulfur compound metabolic process’ (SM), ‘response to oxidative stress’ (OX), and/ or ‘response to metal ion’ (RM) within the list of 38 genes presented in C.

We next examined the overlap between M16 and MP1 and found 65 shared genes, of which 38 were up-regulated under S- at both transcript and protein levels (Fig. 4C). Gene Ontology (GO) enrichment revealed that several of these genes are involved in S compound metabolic process, oxidative stress response and metal ion response (Fig. 4D). Notably, they include two glutathione S-transferases (GSTs), a glutathione reductase, a thioredoxin h, and a cysteine synthase (Fig. 4E). One of the GST was previously identified as a hub in a cluster of antioxidant proteins up-regulated under S- in pea seeds (Henriet et al. 2021). Our results also evidenced *Psat1g179680*, a chloroplastic 5 adenylylsulfate reductase 3, homologous to the Arabidopsis OAS cluster gene *APR3* (*At4g21990*). In total, nine expressed homologs of Arabidopsis OAS cluster genes were identified, all strongly induced under S- and WD/S- and grouped within M16 (Supplementary Fig. S9).

Altogether, these results identified a core set of genes that are tightly co-regulated at both transcript and protein levels under S-, highlighting transcriptional control. The coordinated induction of these genes that include key components of S assimilation and redox homeostasis likely sustains cellular redox balance under low S availability.

### Synergistic gene responses to stress combination

Since WD/S- strongly perturbed phenotypic variables, the transcriptome and the proteome, we next investigated potential synergistic transcriptomic and proteomic responses induced by stress combination. We identified transcripts and proteins that were either specifically up-regulated under WD/S-, or more strongly up-regulated under WD/S- than under either single stress condition (Fig. 5A). This approach does not strictly distinguish additive from synergistic effects; however, given that WD alone affected very few genes, we refer to these responses at synergistic for brevity.

**Fig. 5.**
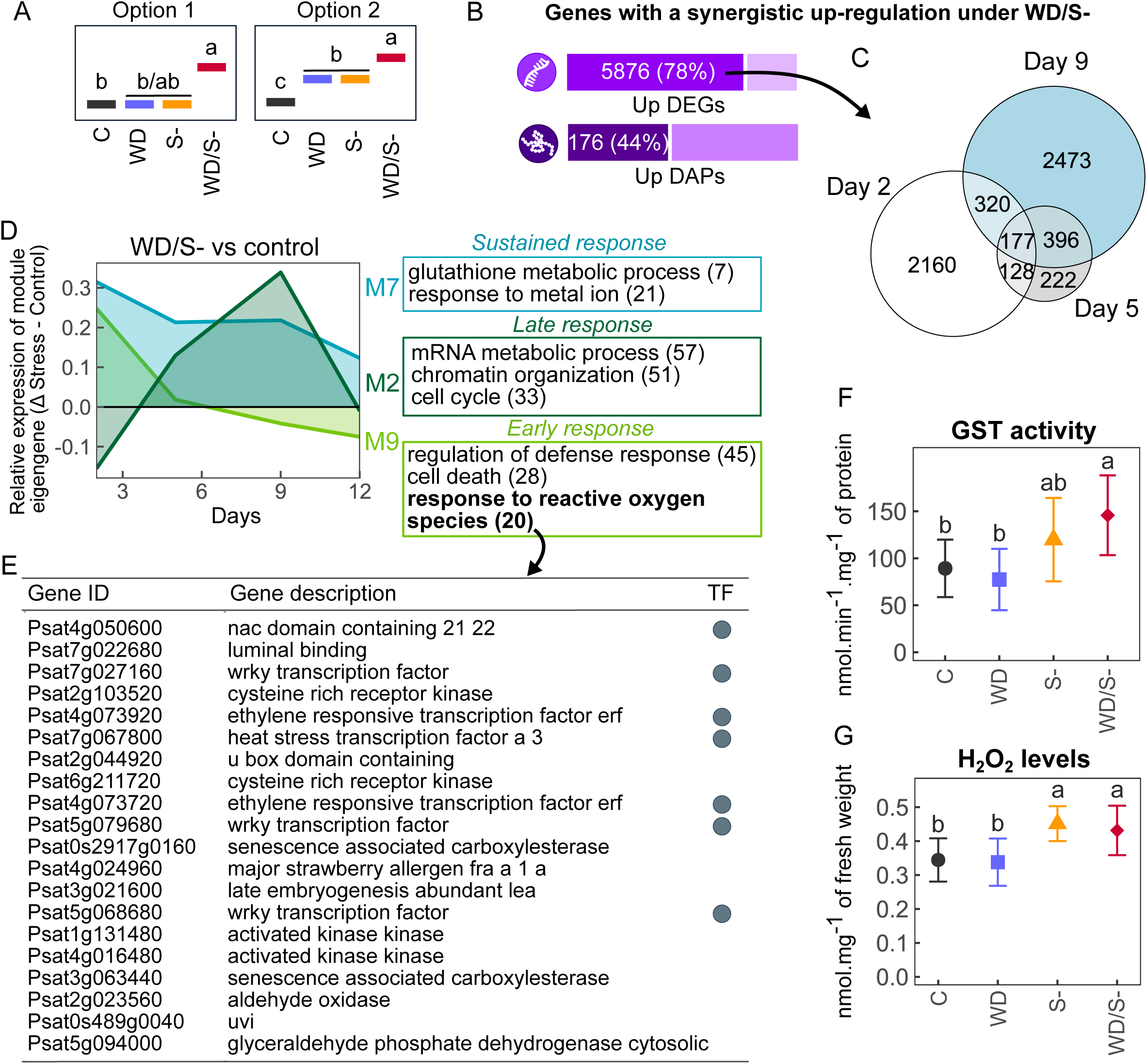
Identification of synergistic responses to stress combination. A, Schematic plot summarizing situations where genes were considered as having a synergistic up-regulation in response to the combined stress as compared to single stress conditions. B, Proportions of differentially expressed genes (DEGs) and differentially accumulated proteins (DAPs) showing a synergistic up-regulation under WD/S-. C, Venn diagram depicting the overlap between lists of genes identified at days 2, 5 and 9 with a synergistic up-regulation under WD/S- at the transcriptome level. D, Area charts representing the average changes in relative expression in response to WD/S- (Δ stress - control), for genes exhibiting a synergistic up-regulation under WD/S- and grouped in modules M2, M7 and M9. Selected enriched biological processes within modules are indicated. Numbers in parentheses indicate gene counts. E, Description of genes with a Gene Ontology annotation corresponding to ‘response to reactive oxygen species’ within the list of genes exhibiting a synergistic up-regulation under WD/S- and grouped in module M9. TF, transcription factor. F and G, Measurement of the glutathione S transferase (GST) activity (F) and quantification of H202 levels (G) in leaves between day 2 and day 9 (means ± S.D). In F and G, two-way ANOVA followed by Tukey’s HSD tests were performed. Different letters indicate significant differences between treatments, determined by Tukey’s HSD tests (P-value < 0.05). GST activity and H202 levels measured at individual days are presented in Supplementary Fig. S10.

In total, 5876 transcripts (78% of up-regulated DEGs) and 176 proteins (44% of up-regulated proteins) displayed synergistic up-regulation under WD/S- (Fig. 5B). Given the higher proportion observed at the transcript level, subsequent analysis focused on the transcriptome. Among the 5876 responsive genes, 2160 (37%) and 2473 (42%) were specific to days 2 and 9, respectively, whereas only 177 were shared across days 2, 5 and 9 (Fig. 5C), consistent with the overall pattern of DEGs under WD/S- (Fig. 3B). The best pea homolog of *AtSLIM1 (Psat5g288560*) showed synergistic up-regulation exclusively at day 9 at the transcript level (Supplementary Dataset S6). Together, these results point to a sequential transcriptional response to WD/S-, with two major waves involving distinct gene sets.

To explore this temporal dynamic functionally, we focused on genes with synergistic up-regulation in modules M2 and M9, representative of the early (M9) and late (M2) responses (Fig. 5D). GO enrichment analysis revealed that the early response involved genes associated with biotic and abiotic stress responses, development, signaling, and ROS response (Fig. 5D; Supplementary Dataset S5). Among the 20 ROS-responsive genes activated early under WD/S-, seven encoded TFs and four encoded kinases (Fig. 5E). The late response was associated with genes involved in mRNA and DNA metabolism, chromatin organization, the cell cycle, and developmental regulation (Fig. 5D).

A third transcriptional signature was identified in module M7, encompassing genes with a sustained response and functions related to metal ion response and glutathione metabolism (Fig. 5D). All seven genes involved in glutathione metabolism encoded GSTs, including one (*Psat3g133720*) also detected by proteomics and showing synergistic protein up-regulation under WD/S- at day 9 (Supplementary Dataset S6).

To assess whether these responses extended to enzyme activity, total GST activity was measured in leaves from day 2 to day 9. Although no significant differences between treatments were observed at individual time points, GST activity was significantly higher under WD/S- over the day 2-day 9 period, confirming a synergistic response at the protein activity level (two-way ANOVA, Fig. 5F, Supplementary Fig. S10).

Quantification of H_2_O_2_ levels between days 2 and 9 showed no significant differences between treatments at individual days; however, a significant increase was observed under both S- and WD/S- (two-way ANOVA) over the day 2-day 9 period (Fig. 5G, Supplementary Fig. S10). Similar magnitudes of response between S- and WD/S- indicated the absence of synergistic response under combined stress.

Collectively, genes identified in modules M7 and M9 may have contributed to mitigating ROS accumulation in leaves, potentially enhancing tolerance to multiple stresses. These results showed that synergistic responses to combined stress span multiple regulatory layers, from transcript accumulation to protein abundance and enzyme activity.

### Protein-specific responses to water deficit and sulfur deficiency

The generally low correlations between transcript and protein abundance patterns (Fig. 4A) suggest that stress responses at these two levels may have occurred independently or with different magnitude or timing. Although the number of differentially accumulated proteins was substantially lower than that of DEGs, we investigated responses that were specific to the protein level. Among the 469 up-regulated proteins, 172 (37%) corresponded to genes not identified as DEGs (Fig. 6A). Of these, 23 were up-regulated across all stress conditions (WD, S- and WD/S-), while 50 were specifically up-regulated under WD/S-. Notably, 37 of these proteins were classified as showing a synergistic response under WD/S- (Fig. 6B and 5B).

**Fig. 6.**
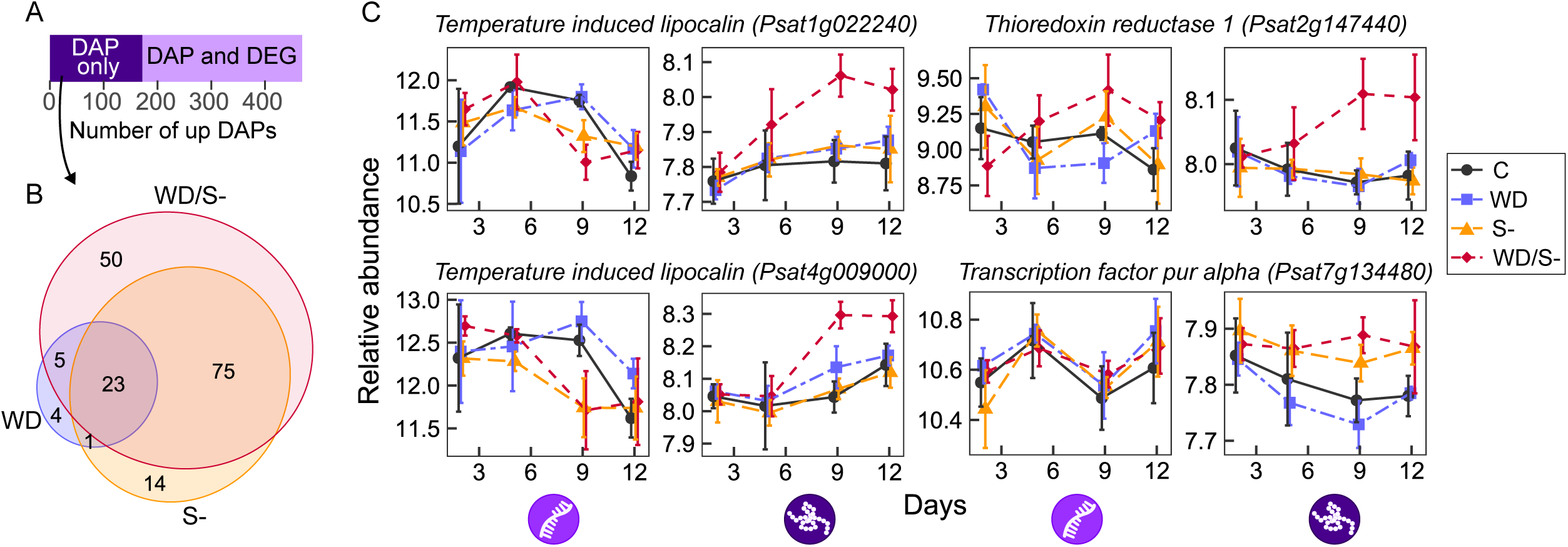
Genes with protein-specific up-regulation in pea leaves under stress. A, Number of differentially accumulated proteins (DAPs) showing higher accumulation in any stress treatment (WD, S- or WD/S-) compared with control, including those with protein-specific up-regulation (DAPs not identified as differentially expressed genes [DEGs]). B, Venn diagram showing overlap of protein-specific up-regulated genes across stress treatments. C, Expression profiles of selected genes, with mRNA levels (left) and protein levels (right) shown (means ± S.D, n = 4 replicates).

No specific GO terms were significantly enriched among these protein-specific genes; however, several encode proteins associated with stress responses. Notably, two corresponded to Temperature Induced Lipocalins (*TILs*, *Psat1g022240* and *Psat4g009000*, Fig. 6C). In Arabidopsis, a single *TIL* gene has been characterized (Frenette Charron et al. 2002, 2005). *Attil* knockout plants are sensitive to temperature and oxidative stress, accumulating elevated levels of ROS, whereas *AtTIL* over-expression enhances tolerance to light stress, freezing, and paraquat treatments (Charron et al. 2008; Chi et al. 2009). In pea, in addition to *Psat1g022240* and *Psat4g009000,* two other neighboring TIL homologs were identified*: Psat4g008960* and *Psat4g009200*. *Psat4g009200* was not expressed and no corresponding protein was detected, whereas *Psat4g008960* displayed an accumulation pattern similar to the other two TILs (Supplementary Fig. S11). Unlike *Psat1g022240* and *Psat4g009000, Psat4g008960* was identified as a DEG and was therefore not included in Fig. 6B. Other genes with protein-specific responses to WD/S- included a thioredoxin reductase 1 (*Psat2g147440*) and a ‘transcription factor pur alpha’ (*Psat7g134480*).

Collectively, these findings reveal a subset of proteins that accumulate under stress independently of transcriptional regulation, pointing to post-transcriptional of translational mechanisms as additional layers of stress response. Among these, TILs may contribute to combined abiotic stress tolerance through mitigation of oxidative damage.

## Discussion

Stress combination can lead to unique molecular responses in plants, which could not be predicted from the effects observed under each individual stress. Here, we focused on the responses of pea to S- and WD, and investigated how the combination of stresses could accentuate the effects of single stresses or induce specific molecular signatures.

### Elemental imbalance and oxidative stress under sulfur deficiency and water deficit

Under S-, applied individually or combined with WD, a strong decrease in leaf S content was observed, together with reduced accumulation of several macro-(phosphorus, potassium, calcium, magnesium) and micro-nutrients (iron), indicating that the uptake and/or transport of elements have been disrupted by S-. Both S- and WD/S- treatments were also characterized by elevated molybdenum contents in leaves, as previously reported in crops exposed to S-limiting conditions (Shinmachi et al. 2010; Maillard et al. 2016). Our results highlighted *PsSULTR4* (*Psat3g185920) and a SULTR* of group 1 (*Psat2g074400*) that were up-regulated under S- and WD/S-, that may play an important role in supplying sulfate under low leaf S status. Under S-, PsSULTR4 drives S distribution from source tissues to seeds to maintain seed production (Bachelet et al. 2024). Molybdate can also be transported by SULTRs (Alhendawi et al. 2005; Fitzpatrick et al. 2008). None of the three annotated pea molybdate transporters (*Psat5g026520*, *Psat6g031280* and *Psat4g172720*) were induced under S- or WD/S-, suggesting that the increase in leaf molybdenum likely results from unspecific transport through up-regulated SULTRs. Similar observations were made in *Brassica napus*, where S- enhanced molybdenum accumulation in leaves and induced SULTR of group 1 in roots, while molybdate transporters showed limited or no transcriptional response (Maillard et al. 2016).

Perturbations of nutrient uptake and transport under S- and WD/S- may also be responsible for the increased accumulation of several metals in leaves, such as cadmium under S-, and iron, zinc, cobalt and arsenic under WD/S-. The increased concentrations of cadmium may result from the activation of transporters of essential or beneficial nutrients (Zhao et al. 2022). For instance, cadmium entry can be mediated by Mn transporters NRAMPs (Cailliatte et al. 2010; Ishimaru et al. 2012; Sasaki et al. 2012) and by the Fe transporter IRT1 (Eide et al. 1996; Korshunova et al. 1999; Connolly et al. 2002). Although our transcriptomics and proteomics data did not reveal significant induction of these specific transporters under the stress treatments applied here, regulation at the post-translational level may still contribute to altered metal uptake. This hypothesis remains speculative and warrants dedicated future investigation.

Consistent with the increased accumulation of metals in leaves, genes involved in metal ion and oxidative stress responses were induced under S- and WD/S-. Connections between redox regulation and metal homeostasis have been well described (for reviews, see Sharma and Dietz, 2009; Ravet and Pilon, 2013). S status plays a central role in this interplay because thiol-containing molecules (glutathione, phytochelatins, cysteine-rich proteins) are ligands for metal buffering and detoxification (Mansoor et al. 2023). This is supported by our results, which include the induction under S- treatments of genes encoding GSTs, a selenium-binding protein and a cysteine synthase, all acting at the interface between S metabolism, metal homeostasis and response to oxidative stress.

Oxidative stress is often associated with increased ROS levels. We identified a set of 20 genes specifically activated under WD/S- at day 2 and involved in response to ROS, despite the absence of similar patterns for leaf H_2_O_2_ levels. This may suggest that ROS-dependent signaling may be activated under WD/S- without a measurable rise in whole-tissue H_2_O_2_, possibly due to localized or transient ROS burst (Mittler et al. 2022). Alternatively, we proposed that these genes may help maintain redox homeostasis under combined stresses, thereby limiting ROS accumulation.

Overall, our results suggest that under S- treatments, disturbances in the uptake and transport of essential and non-essential elements contribute to redox imbalance through ROS-related oxidative stress, amplified by the reduced synthesis of S-containing antioxidant molecules, together with metal-induced redox perturbation.

### Coordinated transcript-protein regulation under stress

Comparison of the transcriptome and proteome datasets revealed a subset of genes that were strongly induced under S- at both the mRNA and protein levels, indicating predominant transcriptional regulation for this group. These genes included GSTs, a cysteine synthase, a thioredoxin, a homolog of *atAPR3*, and other genes associated with sulfur metabolism, oxidative stress and metal ion responses. They were already highly expressed at day 0 under S- treatments, when reduced leaf S contents were detected, and showed no further temporal modulation. This constitutive induction likely reflects an early response to S- that occurred before the first sampling time point, which may also explain the higher correlation between transcript and protein patterns for these genes.

In contrast, global transcript and protein patterns were poorly correlated for other genes, consistent with previous observations. In Maize, for example, co-expression networks of transcripts and proteins built from a developmental atlas were highly distinct, with few shared hubs between networks, the majority of relationships between genes being specific to each network (Walley et al. 2016). The limited correlation between transcript and protein patterns can be attributed to multiple mechanisms. These mechanisms include different stability between transcripts and proteins: mRNA half-lives in plants range from hours on average (Gutiérrez et al. 1999; Narsai et al. 2007), whereas the average half-life of proteins is of several days (Scheurwater et al. 2000; Ishihara et al. 2015; Belouah et al. 2019). As a consequence, protein levels change more gradually than transcript levels, as reported in developing tomato fruits (Belouah et al. 2019).

One third of the differentially accumulated proteins under WD and/or S- were encoded by genes that did not show transcriptional responses. Such protein-specific regulation may indicate selective translation. Under heat stress, for instance, translation of specific mRNAs is prioritized while polysome abundance decreases and global protein synthesis is reduced (Yángüez et al. 2013; Merret et al. 2015; Bonnot and Nagel 2021). Genes with specific responses at the protein level such as TILs may be prioritized for translation under multi-stress conditions, and likely contribute to plant stress tolerance.

### Combined sulfur deficiency and water deficit trigger unique and temporal molecular responses in pea leaves

When occurring alone, WD had a moderate effect on the measured phenotypic variables, and fewer mRNAs and proteins were differentially accumulated in leaves under this treatment as compared to others. However, data obtained at maturity on plants grown in the same experiment showed that the number of reproductive nodes and the one-seed weight were significantly reduced (Henriet et al. 2019). Hence, plants responded to WD over the long term while still acclimating. When WD was combined with S-, synergistic effects were observed at the transcriptome and proteome levels. In facts, 29-fold and 2.6-fold more mRNAs were differentially accumulated under combined stress compared with individual WD and S-, respectively. Such differences were not observed in pea seeds (Henriet et al. 2019, 2021), indicating that transcriptional and translational reprogramming under combined stress mostly happened in leaves.

This study revealed genes with a sustained synergistic up-regulation under combined stress, of which seven encoded GSTs. These may fine-tune redox homeostasis in leaves, presumably resulting from metal (cobalt and arsenate) and ROS accumulation. Many more differentially accumulated transcripts and proteins were observed at days 2 and 9 compared to day 5, indicating that transcriptional and translational responses to combined stress were dynamic in pea leaves. Supported by the up-regulation of genes involved in signaling, response to ROS and developmental processes at day 2, we speculate that the early response may trigger metabolic adjustments, resulting in fewer molecular changes at day 5. These adjustments involve the activation of seven TFs associated with ROS response, which may play a key role in mitigating the negative impact of oxidative stress.

After nine days of combined stress, plants appear to be severely impacted and may activate a “survival” response. The response at day 9 was characterized by the up-regulation of genes involved in chromatin organization and DNA metabolism. Modification of chromatin accessibility may have contributed to an intense transcriptional reprogramming. The best pea homolog of *AtSLIM1 –* a key regulator of S- responses *–* was up-regulated uniquely under combined stress at day 9. In Arabidopsis, *SLIM1* expression is not modulated by S- (Maruyama-Nakashita et al. 2006). Instead, SLIM1 is subjected to constitutive proteasomal degradation, which is repressed at the early stages of S- (Wawrzyńska and Sirko 2020). Our results suggest that when S- co-occur with other stresses, *PsSLIM1* may exhibit transcriptional responses. Further evidence is needed to assess whether a switch in the level of regulation of *PsSLIM1* may occur under multi-stress conditions.

## Conclusion

Although additive effects are observed under stress combination, synergistic responses frequently occur and reveal unpredictable outcomes, as demonstrated here in pea exposed to combined S- and WD. At the mechanistic level, our results evidenced coordinated transcriptional and post-transcriptional responses under combined stress, including the early activation of ROS-responsive TFs and kinases, the sustained induction of GSTs contributing to redox homeostasis, and the accumulation of stress-protective proteins such as TILs that are not transcriptionally regulated. The occurrence of synergistic responses across multiple regulatory layers – from transcriptional reprogramming to translational and post-translational control – highlights the complexity of combined stress responses. Proteins that accumulate specifically under combined S- and WD independently of transcriptional regulation may represent promising targets for improving crop resilience to the stress combinations increasingly associated with climate change.

## Acknowledgments

We thank the members of the 4PMI Platform (Phenotyping Platform for Plant and Plant Microorganisms Interactions, INRAE, Dijon) for their excellent technical support during plant growth and all members of the FILEAS team for helping with sample collection and measurements of plants phenotypic variables. We also thank Mickaël Lamboeuf for the development of the custom image processing algorithm used for the calculation of the leaf surface area, Sylvie Girodet for CN measurements and the GISMO platform (Université de Bourgogne Franche-Comté, Dijon, France) for S measurements; Rémy-Félix Serre from the GeT-PlaGe core facility (Castanet-Tolosan, France) for the sequencing of mRNAs and pre-processing of the mRNA-Seq data. We also thank Marion Prudent for her help with the experimental setup, and Christine Le Signor for helpful discussions to interpret the data.

## Author contribution

Research design: CH, VV and KG; Data acquisition: CH, DA, NR, T Balliau, CB, OL, AO and MZ; Data processing and statistical analyses: T Bonnot, CH, MT, JK and OL; Data exploration: T Bonnot, CH and MT, Data visualization: T Bonnot; Data interpretation: T Bonnot, CH, MT, OL, VV and KG; Writing – original draft: T Bonnot, CH, OL, VV and KG; Manuscript review and editing: all authors.

## Data availability

Raw mass spectrometry files have been deposited to the ProteomeXchange Consortium via the PRIDE partner repository with the dataset identifier PXD048279. Raw RNA-Seq data (fasta files and read counting data) are accessible from the Gene Expression Omnibus (GEO) database, www.ncbi.nlm.nih.gov/geo, with the identifier GSE252351. All other data are available as Supplementary Datasets.

For the purpose of open access, the authors have applied a CC-BY public copyright licence to any Author Accepted Manuscript (AAM) version arising from this submission.

## References

Aarabi F, Kusajima M, Tohge T, Konishi T, Gigolashvili T, Takamune M, Sasazaki Y, Watanabe M, Nakashita H, Fernie AR, et al. Sulfur deficiency-induced repressor proteins optimize glucosinolate biosynthesis in plants. Science Advances. 2016:2(10):e1601087–e1601087. 10.1126/sciadv.1601087

Aarabi F, Rakpenthai A, Barahimipour R, Gorka M, Alseekh S, Zhang Y, Salem MA, Brückner F, Omranian N, Watanabe M, et al. Sulfur deficiency-induced genes affect seed protein accumulation and composition under sulfate deprivation. Plant Physiology. 2021:187(4):2419–2434. 10.1093/plphys/kiab386

Alhendawi RA, Kirkby EA, and Pilbeam DJ. Evidence That Sulfur Deficiency Enhances Molybdenum Transport in Xylem Sap of Tomato Plants. Journal of Plant Nutrition. 2005:28(8):1347–1353. 10.1081/PLN-200067449

Allahham A, Kanno S, Zhang L, and Maruyama-Nakashita A. Sulfur Deficiency Increases Phosphate Accumulation, Uptake, and Transport in Arabidopsis thaliana. International Journal of Molecular Sciences. 2020:21(8):2971. 10.3390/ijms21082971

Allen WM, Berrett S, and Patterson DSP. A biochemical study of experimental Johne’s disease. Journal of Comparative Pathology. 1974:84(3):385–389. 10.1016/0021-9975(74)90013-9

Apodiakou A and Hoefgen R. New insights into the regulation of plant metabolism by O -acetylserine: sulfate and beyond. Journal of Experimental Botany. 2023:74(11):3361–3378. 10.1093/jxb/erad124

Azodi CB, Lloyd JP, and Shiu S-H. The cis-regulatory codes of response to combined heat and drought stress in Arabidopsis thaliana. NAR Genomics and Bioinformatics. 2020:2(3):1–16. 10.1093/nargab/lqaa049

Bachelet F, Sanchez M, Aimé D, Naudé F, Rossin N, Ourry A, Deulvot C, Le Signor C, Vernoud V, Neiers F, et al. The vacuolar sulfate transporter PsSULTR4 is a key determinant of seed yield and protein composition in pea. The Plant Journal. 2024:119(6):2919–2936. 10.1111/tpj.16961

Belouah I, Nazaret C, Pétriacq P, Prigent S, Bénard C, Mengin V, Blein-Nicolas M, Denton AK, Balliau T, Augé S, et al. Modeling Protein Destiny in Developing Fruit. Plant Physiol. 2019:180(3):1709–1724. 10.1104/pp.19.00086

Benjamini Y and Hochberg Y. Controlling the false discovery rate: a practical and powerful approach to multiple testing. Journal of the Royal Statistical Society Series B (Methodological). 1995:57:289–300. 10.2307/2346101

Bolger AM, Lohse M, and Usadel B. Trimmomatic: a flexible trimmer for Illumina sequence data. Bioinformatics. 2014:30(15):2114–2120. 10.1093/bioinformatics/btu170

Bonnot T, Bachelet F, Boudet J, Le Signor C, Bancel E, Vernoud V, Ravel C, and Gallardo K. Sulfur in determining seed protein composition: present understanding of its interaction with abiotic stresses and future directions. Journal of Experimental Botany. 2023:74(11):3276–3285. 10.1093/jxb/erad098

Bonnot T, Gillard M, and Nagel D. A Simple Protocol for Informative Visualization of Enriched Gene Ontology Terms. Bio-101. 2019:e3429. 10.21769/BioProtoc.3429

Bonnot T, Martre P, Hatte V, Dardevet M, Leroy P, Bénard C, Falagán N, Martin-Magniette M-L, Deborde C, Moing A, et al. Omics Data Reveal Putative Regulators of Einkorn Grain Protein Composition under Sulfur Deficiency. Plant Physiology. 2020:183(2):501–516. 10.1104/pp.19.00842

Bonnot T and Nagel DH. Time of the day prioritizes the pool of translating mRNAs in response to heat stress. The Plant Cell. 2021:33(7):2164–2182. 10.1093/plcell/koab113

Borja Reis AF de, Rosso LHM, Davidson D, Kovács P, Purcell LC, Below FE, Casteel SN, Knott C, Kandel H, Naeve SL, et al. Sulfur fertilization in soybean: A meta-analysis on yield and seed composition. European Journal of Agronomy. 2021:127:126285. 10.1016/j.eja.2021.126285

Cailliatte R, Schikora A, Briat J-F, Mari S, and Curie C. High-Affinity Manganese Uptake by the Metal Transporter NRAMP1 Is Essential for Arabidopsis Growth in Low Manganese Conditions. The Plant Cell. 2010:22(3):904–917. 10.1105/tpc.109.073023

Carrillo R, Moreno I, Romero LC, Aroca A, and Gotor C. Hydrogen sulfide-induced barley resilience to drought and salinity through protein persulfidation. Plant Physiology and Biochemistry. 2025:221:109644. 10.1016/j.plaphy.2025.109644

Charron J-BF, Ouellet F, Houde M, and Sarhan F. The plant Apolipoprotein D ortholog protects Arabidopsis against oxidative stress. BMC Plant Biology. 2008:8(1):86. 10.1186/1471-2229-8-86

Chi W-T, Fung RWM, Liu H-C, Hsu C-C, and Charng Y-Y. Temperature-induced lipocalin is required for basal and acquired thermotolerance in Arabidopsis. Plant, Cell & Environment. 2009:32(7):917–927. 10.1111/j.1365-3040.2009.01972.x

Cohen I, Zandalinas SI, Huck C, Fritschi FB, and Mittler R. Meta-analysis of drought and heat stress combination impact on crop yield and yield components. Physiologia Plantarum. 2021:171(1):66–76. 10.1111/ppl.13203

Connolly EL, Fett JP, and Guerinot ML. Expression of the IRT1 Metal Transporter Is Controlled by Metals at the Levels of Transcript and Protein Accumulation. Plant Cell. 2002:14(6):1347–1357. 10.1105/tpc.001263

Craig R and Beavis RC. TANDEM: matching proteins with tandem mass spectra. Bioinformatics. 2004:20(9):1466–1467. 10.1093/bioinformatics/bth092

Czékus Z, Farkas M, Bakacsy L, Ördög A, Gallé Á, and Poór P. Time-Dependent Effects of Bentazon Application on the Key Antioxidant Enzymes of Soybean and Common Ragweed. Sustainability. 2020:12(9):3872. 10.3390/su12093872

Eide D, Broderius M, Fett J, and Guerinot ML. A novel iron-regulated metal transporter from plants identified by functional expression in yeast. Proceedings of the National Academy of Sciences. 1996:93(11):5624–5628. 10.1073/pnas.93.11.5624

Fitzpatrick KL, Tyerman SD, and Kaiser BN. Molybdate transport through the plant sulfate transporter SHST1. FEBS Letters. 2008:582(10):1508–1513. 10.1016/j.febslet.2008.03.045

Frenette Charron J-B, Breton G, Badawi M, and Sarhan F. Molecular and structural analyses of a novel temperature stress-induced lipocalin from wheat and Arabidopsis. FEBS Letters. 2002:517(1–3):129–132. 10.1016/S0014-5793(02)02606-6

Frenette Charron J-B, Ouellet F, Pelletier M, Danyluk J, Chauve C, and Sarhan F. Identification, Expression, and Evolutionary Analyses of Plant Lipocalins. Plant Physiol. 2005:139(4):2017–2028. 10.1104/pp.105.070466

Gutiérrez RA, MacIntosh GC, and Green PJ. Current perspectives on mRNA stability in plants: multiple levels and mechanisms of control. Trends in Plant Science. 1999:4(11):429–438. 10.1016/S1360-1385(99)01484-3

Haneklaus S, Bloem E, and Schnug E. History of Sulfur Deficiency in Crops. In. Sulfur: A Missing Link between Soils, Crops, and Nutrition. (John Wiley & Sons, Ltd), pp. 45–58–6. 10.2134/agronmonogr50.c4

Henriet C, Aimé D, Térézol M, Kilandamoko A, Rossin N, Combes-Soia L, Labas V, Serre R-F, Prudent M, Kreplak J, et al. Water stress combined with sulfur deficiency in pea affects yield components but mitigates the effect of deficiency on seed globulin composition. Journal of Experimental Botany. 2019:70(16):4287–4304. 10.1093/jxb/erz114

Henriet C, Balliau T, Aimé D, Le Signor C, Kreplak J, Zivy M, Gallardo K, and Vernoud V. Proteomics of developing pea seeds reveals a complex antioxidant network underlying the response to sulfur deficiency and water stress. Journal of Experimental Botany. 2021:72(7):2611–2626. 10.1093/jxb/eraa571

Hubberten H-M, Klie S, Caldana C, Degenkolbe T, Willmitzer L, and Hoefgen R. Additional role of O-acetylserine as a sulfur status-independent regulator during plant growth. The Plant Journal. 2012:70(4):666–677. 10.1111/j.1365-313X.2012.04905.x

IPCC. IPCC, 2023: Climate Change 2023: Synthesis Report. A Report of the Intergovernmental Panel on Climate Change. Contribution of Working Groups I, II and III to the Sixth Assessment Report of the Intergovernmental Panel on Climate Change (Geneva, Switzerland).

Ishihara H, Obata T, Sulpice R, Fernie AR, and Stitt M. Quantifying Protein Synthesis and Degradation in Arabidopsis by Dynamic 13CO2 Labeling and Analysis of Enrichment in Individual Amino Acids in Their Free Pools and in Protein. Plant Physiol. 2015:168(1):74–93. 10.1104/pp.15.00209

Ishimaru Y, Takahashi R, Bashir K, Shimo H, Senoura T, Sugimoto K, Ono K, Yano M, Ishikawa S, Arao T, et al. Characterizing the role of rice NRAMP5 in Manganese, Iron and Cadmium Transport. Scientific Reports. 2012:2(1):286. 10.1038/srep00286

Jacques C, Forest M, Durey V, Salon C, Ourry A, and Prudent M. Transient Nutrient Deficiencies in Pea: Consequences on Nutrient Uptake, Remobilization, and Seed Quality. Front Plant Sci. 2021:12. 10.3389/fpls.2021.785221

Jurado-Flores A, Aroca A, Romero LC, and Gotor C. Sulfide promotes tolerance to drought through protein persulfidation in Arabidopsis. J Exp Bot. 2023:74(15):4654–4669. 10.1093/jxb/erad165

Kim D, Langmead B, and Salzberg SL. HISAT: a fast spliced aligner with low memory requirements. Nature Methods. 2015:12(4):357–360. 10.1038/nmeth.3317

Kolde R. pheatmap: Pretty Heatmaps. 2019.

Korshunova YO, Eide D, Gregg Clark W, Lou Guerinot M, and Pakrasi HB. The IRT1 protein from Arabidopsis thaliana is a metal transporter with a broad substrate range. Plant Mol Biol. 1999:40(1):37–44. 10.1023/A:1026438615520

Kreplak J, Madoui M-A, Cápal P, Novák P, Labadie K, Aubert G, Bayer PE, Gali KK, Syme RA, Main D, et al. A reference genome for pea provides insight into legume genome evolution. Nature Genetics. 2019:51(9):1411–1422. 10.1038/s41588-019-0480-1

Langella O, Valot B, Balliau T, Blein-Nicolas M, Bonhomme L, and Zivy M. X!TandemPipeline: A Tool to Manage Sequence Redundancy for Protein Inference and Phosphosite Identification. Journal of Proteome Research. 2017:16(2):494–503. 10.1021/acs.jproteome.6b00632

Langfelder P and Horvath S. WGCNA: an R package for weighted correlation network analysis. BMC Bioinformatics. 2008:9(1):559. 10.1186/1471-2105-9-559

Larsson J. eulerr: Area-Proportional Euler and Venn Diagrams with Ellipses. 2022.

Lecomte C, Richer V, Gauffreteau A, Jeuffroy MH, Bouviala M, Brun C, Buridan C, Klein A, Lantoine FX, Marchand D, et al. Combining a multi-environment trial and a diagnosis method to assess potential yield and main limiting factors of three highly different pea types. European Journal of Agronomy. 2023:146(April). 10.1016/j.eja.2023.126823

Li L-H, Yi H-L, Xiu-Ping Liu, and Qi H-X. Sulfur dioxide enhance drought tolerance of wheat seedlings through H2S signaling. Ecotoxicology and Environmental Safety. 2021:207:111248. 10.1016/j.ecoenv.2020.111248

Liao Y, Smyth GK, and Shi W. featureCounts: an efficient general purpose program for assigning sequence reads to genomic features. Bioinformatics. 2014:30(7):923–930. 10.1093/bioinformatics/btt656

Love MI, Huber W, and Anders S. Moderated estimation of fold change and dispersion for RNA-seq data with DESeq2. Genome Biology. 2014:15(12):550. 10.1186/s13059-014-0550-8

Mahalingam R, Duhan N, Kaundal R, Smertenko A, Nazarov T, and Bregitzer P. Heat and drought induced transcriptomic changes in barley varieties with contrasting stress response phenotypes. Frontiers in Plant Science. 2022:13(December):1–24. 10.3389/fpls.2022.1066421

Maillard A, Etienne P, Diquélou S, Trouverie J, Billard V, Yvin J-C, and Ourry A. Nutrient deficiencies modify the ionomic composition of plant tissues: a focus on cross-talk between molybdenum and other nutrients in Brassica napus. Journal of Experimental Botany. 2016:67(19):5631–5641. 10.1093/jxb/erw322

Mansoor S, Ali A, Kour N, Bornhorst J, AlHarbi K, Rinklebe J, Abd El Moneim D, Ahmad P, and Chung YS. Heavy Metal Induced Oxidative Stress Mitigation and ROS Scavenging in Plants. Plants (Basel). 2023:12(16):3003. 10.3390/plants12163003

Maruyama-Nakashita A, Nakamura Y, Tohge T, Saito K, and Takahashi H. Arabidopsis SLIM1 Is a Central Transcriptional Regulator of Plant Sulfur Response and Metabolism. The Plant Cell. 2006:18(11):3235–3251. 10.1105/tpc.106.046458

Merret R, Nagarajan VK, Carpentier M-C, Park S, Favory J-J, Descombin J, Picart C, Charng Y, Green PJ, Deragon J-M, et al. Heat-induced ribosome pausing triggers mRNA co-translational decay in *Arabidopsis thaliana*. Nucleic Acids Research. 2015:43(8):4121–4132. 10.1093/nar/gkv234

Mittler R. Abiotic stress, the field environment and stress combination. Trends in Plant Science. 2006:11(1):15–19. 10.1016/j.tplants.2005.11.002

Mittler R, Zandalinas SI, Fichman Y, and Van Breusegem F. Reactive oxygen species signalling in plant stress responses. Nat Rev Mol Cell Biol. 2022:23(10):663–679. 10.1038/s41580-022-00499-2

Mondal S, Pramanik K, Panda D, Dutta D, Karmakar S, and Bose B. Sulfur in Seeds: An Overview. Plants. 2022:11(3):450. 10.3390/plants11030450

Narsai R, Howell KA, Millar AH, O’Toole N, Small I, and Whelan J. Genome-Wide Analysis of mRNA Decay Rates and Their Determinants in Arabidopsis thaliana. Plant Cell. 2007:19(11):3418–3436. 10.1105/tpc.107.055046

Noctor G, Mhamdi A, and Foyer CH. Oxidative stress and antioxidative systems: recipes for successful data collection and interpretation. Plant, Cell & Environment. 2016:39(5):1140–1160. 10.1111/pce.12726

Paine RT, Tegner MJ, and Johnson E a. Ecological Surprises. Ecosystems. 1998:1(July):535–545.

Poisson E, Trouverie J, Brunel-Muguet S, Akmouche Y, Pontet C, Pinochet X, and Avice J-C. Seed Yield Components and Seed Quality of Oilseed Rape Are Impacted by Sulfur Fertilization and Its Interactions With Nitrogen Fertilization. Frontiers in Plant Science. 2019:10. 10.3389/fpls.2019.00458

R Core Team. R: A Language and environment for statistical computing. 2022.

Rasmussen S, Barah P, Suarez-Rodriguez MC, Bressendorff S, Friis P, Costantino P, Bones AM, Nielsen HB, and Mundy J. Transcriptome responses to combinations of stresses in *Arabidopsis*. Plant Physiology. 2013:161(4):1783–1794. 10.1104/pp.112.210773

Ravet K and Pilon M. Copper and Iron Homeostasis in Plants: The Challenges of Oxidative Stress. Antioxid Redox Signal. 2013:19(9):919–932. 10.1089/ars.2012.5084

Ristova D and Kopriva S. Sulfur signaling and starvation response in Arabidopsis. iScience. 2022:25(5):104242. 10.1016/j.isci.2022.104242

Rizhsky L, Liang H, and Mittler R. The Combined Effect of Drought Stress and Heat Shock on Gene Expression in Tobacco. Plant Physiology. 2002:130(3):1143–1151. 10.1104/pp.006858

Rizhsky L, Liang H, Shuman J, Shulaev V, Davletova S, and Mittler R. When defense pathways collide. The response of arabidopsis to a combination of drought and heat stress 1[w]. Plant Physiology. 2004:134(4):1683–1696. 10.1104/pp.103.033431

Samanta S, Singh A, and Roychoudhury A. Involvement of Sulfur in the Regulation of Abiotic Stress Tolerance in Plants. In. Protective Chemical Agents in the Amelioration of Plant Abiotic Stress. (Wiley), pp. 437–466. 10.1002/9781119552154.ch22

Sasaki A, Yamaji N, Yokosho K, and Ma JF. Nramp5 Is a Major Transporter Responsible for Manganese and Cadmium Uptake in Rice. The Plant Cell. 2012:24(5):2155–2167. 10.1105/tpc.112.096925

Scherer HW. Sulphur in crop production. European Journal of Agronomy. 2001:14(2):81–111. 10.1016/S1161-0301(00)00082-4

Scheurwater I, Dünnebacke M, Eising R, and Lambers H. Respiratory costs and rate of protein turnover in the roots of a fast-growing ( Dactylis glomerata L.) and a slow-growing ( Festuca ovina L.) grass species. J Exp Bot. 2000:51(347):1089–1097. 10.1093/jxb/51.347.1089

Schmittgen TD and Livak KJ. Analyzing real-time PCR data by the comparative CT method. Nat Protoc. 2008:3(6):1101–1108. 10.1038/nprot.2008.73

Sharma SS and Dietz K-J. The relationship between metal toxicity and cellular redox imbalance. Trends in Plant Science. 2009:14(1):43–50. 10.1016/j.tplants.2008.10.007

Shinmachi F, Buchner P, Stroud JL, Parmar S, Zhao F-J, McGrath SP, and Hawkesford MJ. Influence of Sulfur Deficiency on the Expression of Specific Sulfate Transporters and the Distribution of Sulfur, Selenium, and Molybdenum in Wheat. Plant Physiology. 2010:153(1):327–336. 10.1104/pp.110.153759

Smoot ME, Ono K, Ruscheinski J, Wang P-L, and Ideker T. Cytoscape 2.8: new features for data integration and network visualization. Bioinformatics. 2011:27(3):431–432. 10.1093/bioinformatics/btq675

Takahashi H, Watanabe-Takahashi A, Smith FW, Blake-Kalff M, Hawkesford MJ, and Saito K. The roles of three functional sulphate transporters involved in uptake and translocation of sulphate in Arabidopsis thaliana. The Plant Journal. 2000:23(2):171–182. 10.1046/j.1365-313x.2000.00768.x

Tan JW, Shinde H, Tesfamicael K, Hu Y, Fruzangohar M, Tricker P, Baumann U, Edwards EJ, and Rodríguez López CM. Global transcriptome and gene co-expression network analyses reveal regulatory and non-additive effects of drought and heat stress in grapevine. Frontiers in Plant Science. 2023:14(February):1–15. 10.3389/fpls.2023.1096225

Valot B, Langella O, Nano E, and Zivy M. MassChroQ: A versatile tool for mass spectrometry quantification. PROTEOMICS. 2011:11(17):3572–3577. 10.1002/pmic.201100120

Vidmar JJ, Tagmount A, Cathala N, Touraine B, and Davidian J-CE. Cloning and characterization of a root specific high-affinity sulfate transporter from *Arabidopsis thaliana*1. FEBS Letters. 2000:475(1):65–69. 10.1016/S0014-5793(00)01615-X

Walley JW, Sartor RC, Shen Z, Schmitz RJ, Wu KJ, Urich MA, Nery JR, Smith LG, Schnable JC, Ecker JosephR, et al. Integration of omic networks in a developmental atlas of maize. Science. 2016:353(6301):814–818. 10.1126/science.aag1125

Wawrzyńska A and Sirko A. Proteasomal Degradation of Proteins Is Important for the Proper Transcriptional Response to Sulfur Deficiency Conditions in Plants. Plant Cell Physiol. 2020:61(9):1548–1564. 10.1093/pcp/pcaa076

Wickham H. ggplot2: Elegant Graphics for Data Analysis. 2016.

Yángüez E, Castro-Sanz AB, Fernández-Bautista N, Oliveros JC, and Castellano MM. Analysis of genome-wide changes in the translatome of *Arabidopsis* seedlings subjected to heat stress. PLoS ONE. 2013:8(8):e71425. 10.1371/journal.pone.0071425

Yoshimoto N, Takahashi H, Smith FW, Yamaya T, and Saito K. Two distinct high-affinity sulfate transporters with different inducibilities mediate uptake of sulfate in *Arabidopsis* roots. The Plant Journal. 2002:29(4):465–473. 10.1046/j.0960-7412.2001.01231.x

Yu G, Wang L-G, Han Y, and He Q-Y. clusterProfiler: an R Package for Comparing Biological Themes Among Gene Clusters. OMICS. 2012:16(5):284–287. 10.1089/omi.2011.0118

Zandalinas SI, Sengupta S, Fritschi FB, Azad RK, Nechushtai R, and Mittler R. The impact of multifactorial stress combination on plant growth and survival. New Phytologist. 2021:230(3):1034–1048. 10.1111/nph.17232

Zhang J, Aroca A, Hervás M, Navarro JA, Moreno I, Xie Y, Romero LC, and Gotor C. Analysis of sulfide signaling in rice highlights specific drought responses. J Exp Bot. 2024:75(16):5130–5145. 10.1093/jxb/erae249

Zhao F-J, Tang Z, Song J-J, Huang X-Y, and Wang P. Toxic metals and metalloids: Uptake, transport, detoxification, phytoremediation, and crop improvement for safer food. Molecular Plant. 2022:15(1):27–44. 10.1016/j.molp.2021.09.016

Zheng Y, Jiao C, Sun H, Rosli HG, Pombo MA, Zhang P, Banf M, Dai X, Martin GB, Giovannoni JJ, et al. iTAK: A Program for Genome-wide Prediction and Classification of Plant Transcription Factors, Transcriptional Regulators, and Protein Kinases. Molecular Plant. 2016:9(12):1667–1670. 10.1016/j.molp.2016.09.014

