## Supplemental Figures for "Combined sulfur deficiency and water deficit trigger synergistic redox adjustments through coordinated transcript-protein regulation in pea"

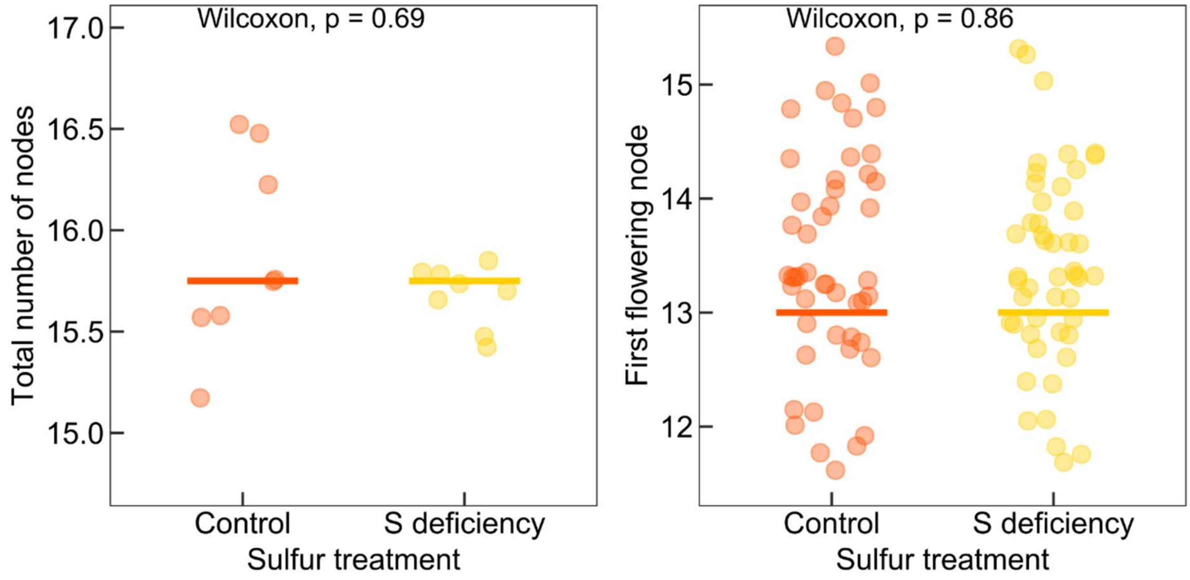


**Supplementary Fig. S1.** Influence of sulfur deficiency treatment on the total number of nodes per plant (left) and on the first flowering node (right). Horizontal lines represent medians of *n* = 8 (left) and *n* = 48 (right) individual plants. Wilcoxon signed-rank tests have been performed to compare medians between groups.

A

| Gene ID | Annotation | Primer F (5'-3') | Primer R (5'-3') |
|---|---|---|---|
| Psat3g037400 | Sulfite exporter TauE/SafE | CTGTTCTAGCTGGATTTGTGGGAC | GACCCCTGAGAGGATGAACAC |
| Psat5g207000 | Eukaryotic protein of unknown function (DUF953) | ACCAACATGGAGGAACCCACAC | GCGACCTGTGACAGCATCATTCC |
| Psat1g179680 | APS reductase | AGGGTCAGGAGTGAAGGTTGGA | GTTTGGGGAAGAAGAGGATTGTG |
| Psat4g082000 | Adenylylsulphate kinase | CCGTGTGGCTGTGAGATAGTC | GTGTCCACTCTTCTCCAAATAGG |
| Psat3g042160 | S-adenosylmethionine synthetase + C-terminal domain | CTGTTCTAGCTGGATTTGTGGGAC | GACCCCTGAGAGGATGAACAC |
| Psat5g091400 | α-Tubulin | TTGGTGCTGAGTCTGGTGATGG | TTGCACAACACAAAAGAACAAGCCC |

B

| | | Day 0 | Day 2 | | | Day 5 | | | Day 9 | | | Day 12 | | |
|---|---|---|---|---|---|---|---|---|---|---|---|---|---|---|
| Technique | ID | S- | WD | S- | WD/S- | WD | S- | WD/S- | WD | S- | WD/S- | WD | S- | WD/S- |
| RNASeq | Psat1g179680 | 1.12 | -0.67 | 1.68 | 1.16 | 0.06 | 0.14 | 0.08 | -0.07 | 1.59 | 1.36 | -0.06 | 1.43 | 1.23 |
| qPCR | Psat1g179680 | 0.15 | -0.93 | 2.31 | 1.43 | -0.17 | -0.03 | 0.01 | 0.93 | 1.44 | 1.83 | 0.73 | 1.95 | 1.01 |
| RNASeq | Psat3g037400 | -2.50 | -0.04 | -3.06 | -3.41 | -0.23 | -1.78 | -1.77 | 0.05 | -2.04 | -1.93 | -0.16 | -2.55 | -2.31 |
| qPCR | Psat3g037400 | -2.25 | 0.57 | -3.51 | -3.33 | -2.85 | -5.98 | -4.16 | 1.27 | -2.63 | -2.68 | -0.49 | -3.11 | -3.15 |
| RNASeq | Psat3g042160 | -2.24 | 0.17 | -1.00 | -0.91 | -0.38 | -2.29 | -1.92 | -1.19 | -2.76 | -2.28 | -0.47 | -2.44 | -1.29 |
| qPCR | Psat3g042160 | -2.00 | -0.19 | -0.68 | -0.57 | 0.31 | -2.13 | -2.09 | -0.94 | -2.34 | -2.61 | 0.36 | -1.20 | -0.66 |
| RNASeq | Psat4g082000 | -1.85 | 0.00 | -1.93 | -1.86 | -0.21 | -1.33 | -1.63 | -0.07 | -2.20 | -2.20 | 0.04 | -2.09 | -2.24 |
| qPCR | Psat4g082000 | -2.82 | -0.46 | -2.57 | -3.06 | 0.78 | -2.06 | -1.61 | -0.24 | -3.03 | -3.44 | 0.07 | -2.28 | -2.56 |
| RNASeq | Psat5g207000 | 3.21 | -0.16 | 2.86 | 3.58 | 1.70 | 3.85 | 2.98 | 0.19 | 3.18 | 3.35 | 0.23 | 3.28 | 3.07 |
| qPCR | Psat5g207000 | 2.78 | -0.82 | 2.91 | 3.00 | -1.34 | 3.16 | 1.22 | 0.43 | 2.62 | 2.66 | -0.10 | 4.23 | 3.75 |

C

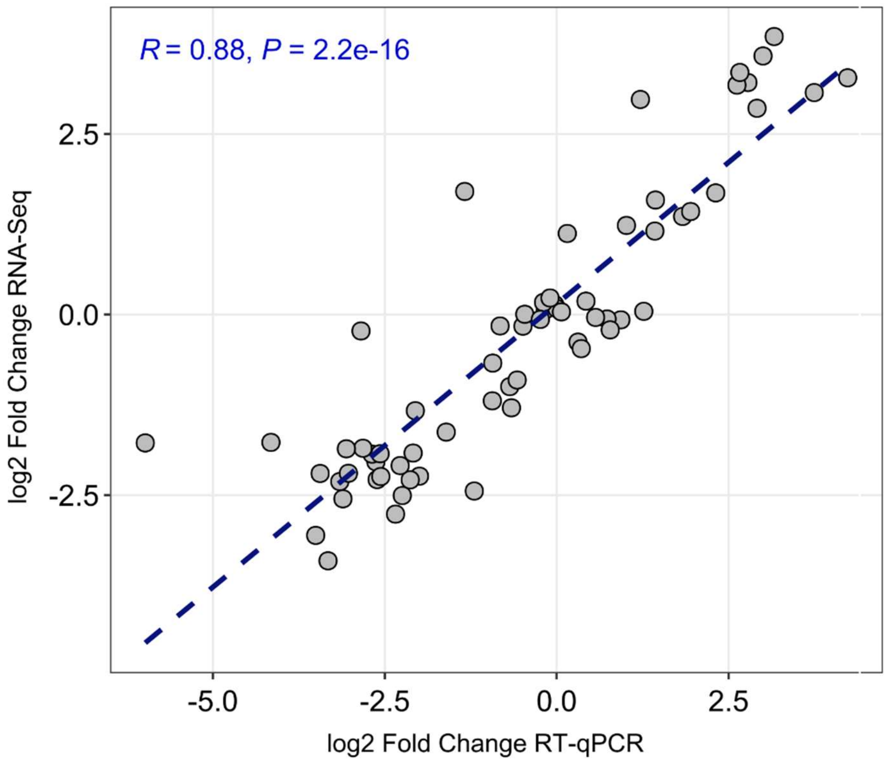


**Supplementary Fig. S2.** RT-qPCR validation of RNA-seq results. Five genes showing contrasted patterns of expression across time and in response to treatments were selected. A, Primer sequences used in RT-qPCR analyses. B, RNA-Seq and RT-qPCR $Log_2$ Fold Change (Stress vs control) values. C, Correlation (Spearman) analysis between RNA-seq and RT-qPCR $Log_2$ Fold Change results from the same RNA samples. WD: water deficit, S-: Sulfur deficiency.

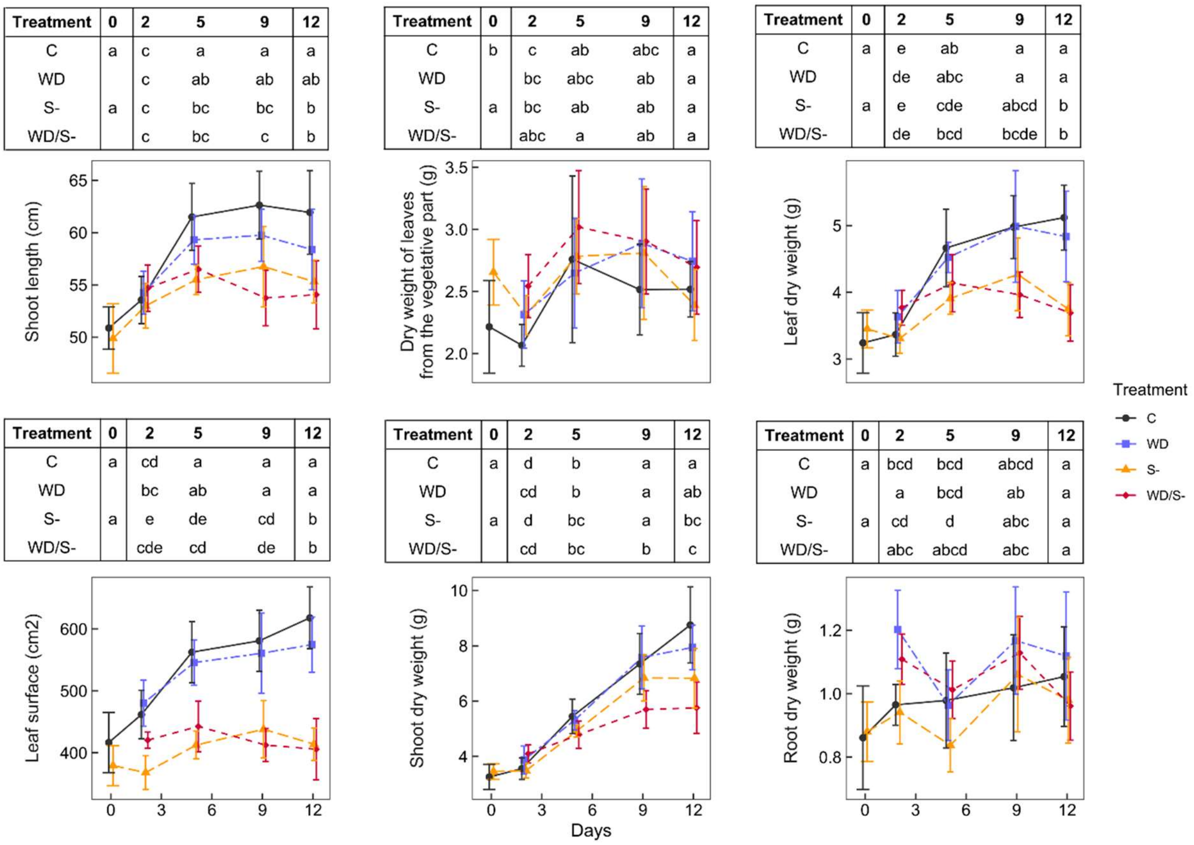


| Treatment | 0 | 2 | 5 | 9 | 12 |
|---|---|---|---|---|---|
| C | a | c | a | a | a |
| WD | | c | ab | ab | ab |
| S- | a | c | bc | bc | b |
| WD/S- | | c | bc | c | b |

| Treatment | 0 | 2 | 5 | 9 | 12 |
|---|---|---|---|---|---|
| C | b | c | ab | abc | a |
| WD | | bc | abc | ab | a |
| S- | a | bc | ab | ab | a |
| WD/S- | | abc | a | ab | a |

| Treatment | 0 | 2 | 5 | 9 | 12 |
|---|---|---|---|---|---|
| C | a | e | ab | a | a |
| WD | | de | abc | a | a |
| S- | a | e | cde | abcd | b |
| WD/S- | | de | bcd | bcde | b |

| Treatment | 0 | 2 | 5 | 9 | 12 |
|---|---|---|---|---|---|
| C | a | cd | a | a | a |
| WD | | bc | ab | a | a |
| S- | a | e | de | cd | b |
| WD/S- | | cde | cd | de | b |

| Treatment | 0 | 2 | 5 | 9 | 12 |
|---|---|---|---|---|---|
| C | a | d | b | a | a |
| WD | | cd | b | a | ab |
| S- | a | d | bc | a | bc |
| WD/S- | | cd | bc | b | c |

| Treatment | 0 | 2 | 5 | 9 | 12 |
|---|---|---|---|---|---|
| C | a | bcd | bcd | abcd | a |
| WD | | a | bcd | ab | a |
| S- | a | cd | d | abc | a |
| WD/S- | | abc | abcd | abc | a |

**Supplementary Fig. S3.** Profiles of phenotypic variables not presented in Figure 1C. Data are means ± S.D. of *n* = 8. Above each plot, different letters represent significant differences between treatments. Three separate statistical analyses were performed: comparison of S- and control at day 0 (Student's t-test); comparisons of all treatments from day 2 to day 9 (two-way ANOVA followed by Tukey's HSD tests); comparisons of all treatments at day 12 during the plant recovery (one-way ANOVA followed by Tukey's HSD tests, see methods for details). Raw data and statistical results are presented in Supplementary Datasets S1 and S2, respectively.

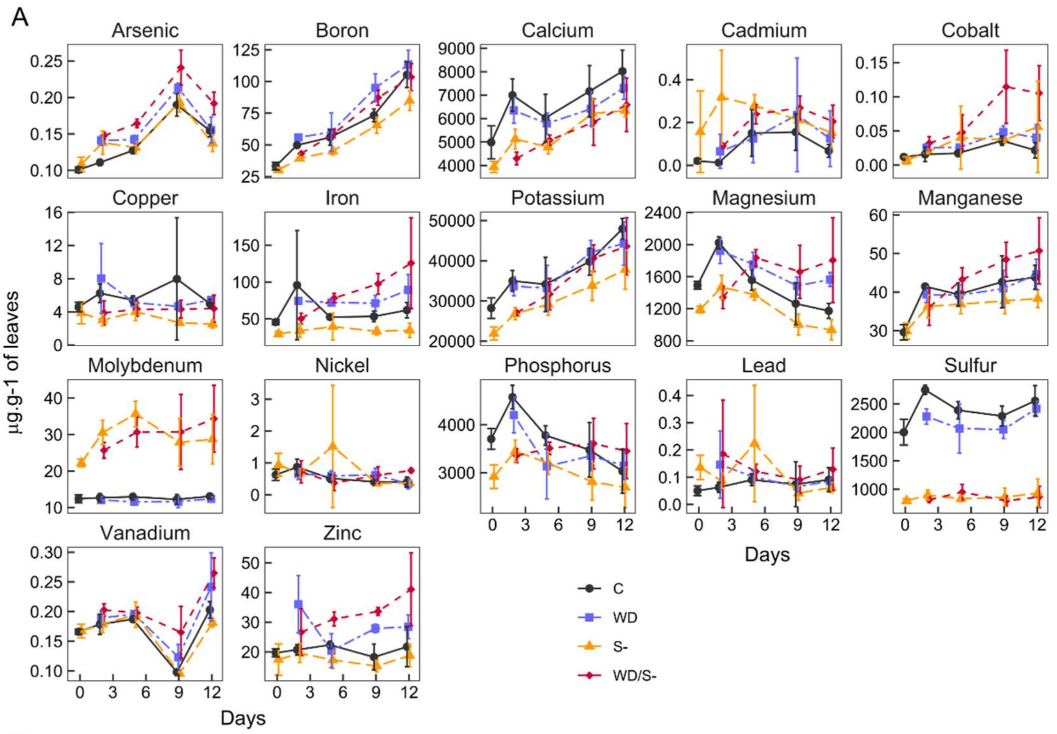


B

**Arsenic**

| Treatment | 0 | 2 | 5 | 9 | 12 |
|---|---|---|---|---|---|
| C | a | f | ef | bc | b |
| WD | | de | de | ab | b |
| S- | a | def | ef | b | b |
| WD/S- | | de | cd | a | a |

**Boron**

| Treatment | 0 | 2 | 5 | 9 | 12 |
|---|---|---|---|---|---|
| C | a | defg | def | bc | ab |
| WD | | def | cde | a | a |
| S- | a | g | efg | cd | b |
| WD/S- | | fg | cdef | ab | ab |

**Calcium**

| Treatment | 0 | 2 | 5 | 9 | 12 |
|---|---|---|---|---|---|
| C | a | a | ab | a | a |
| WD | | ab | abc | ab | ab |
| S- | a | bc | bc | ab | b |
| WD/S- | | c | bc | abc | ab |

**Cadmium**

| Treatment | 0 | 2 | 5 | 9 | 12 |
|---|---|---|---|---|---|
| C | a | b | ab | ab | a |
| WD | | ab | ab | ab | a |
| S- | a | a | ab | ab | a |
| WD/S- | | ab | ab | ab | a |

**Cobalt**

| Treatment | 0 | 2 | 5 | 9 | 12 |
|---|---|---|---|---|---|
| C | | b | b | b | a |
| WD | | b | b | b | a |
| S- | | b | b | b | a |
| WD/S- | | b | b | a | a |

**Copper**

| Treatment | 0 | 2 | 5 | 9 | 12 |
|---|---|---|---|---|---|
| C | a | a | a | a | a |
| WD | | a | a | a | a |
| S- | a | a | a | a | b |
| WD/S- | | a | a | a | a |

**Iron**

| Treatment | 0 | 2 | 5 | 9 | 12 |
|---|---|---|---|---|---|
| C | a | ab | abc | abc | ab |
| WD | | abc | abc | abc | ab |
| S- | b | c | bc | c | b |
| WD/S- | | abc | abc | a | a |

**Potassium**

| Treatment | 0 | 2 | 5 | 9 | 12 |
|---|---|---|---|---|---|
| C | a | abc | abc | ab | a |
| WD | | abc | abc | a | a |
| S- | b | c | c | abc | a |
| WD/S- | | c | bc | ab | a |

**Magnesium**

| Treatment | 0 | 2 | 5 | 9 | 12 |
|---|---|---|---|---|---|
| C | a | a | bcde | ef | bc |
| WD | | ab | abcd | cde | ab |
| S- | b | cde | def | f | c |
| WD/S- | | ef | abc | abcde | a |

**Manganese**

| Treatment | 0 | 2 | 5 | 9 | 12 |
|---|---|---|---|---|---|
| C | a | ab | ab | ab | ab |
| WD | | ab | b | ab | ab |
| S- | a | b | b | b | b |
| WD/S- | | b | ab | a | a |

**Molybdenum**

| Treatment | 0 | 2 | 5 | 9 | 12 |
|---|---|---|---|---|---|
| C | b | b | b | b | b |
| WD | | b | b | b | b |
| S- | a | a | a | a | a |
| WD/S- | | a | a | a | a |

**Nickel**

| Treatment | 0 | 2 | 5 | 9 | 12 |
|---|---|---|---|---|---|
| C | a | a | a | a | b |
| WD | | a | a | a | b |
| S- | a | a | a | a | b |
| WD/S- | | a | a | a | a |

**Phosphorus**

| Treatment | 0 | 2 | 5 | 9 | 12 |
|---|---|---|---|---|---|
| C | a | a | abc | bcd | a |
| WD | | ab | cd | bcd | a |
| S- | b | bcd | cd | d | a |
| WD/S- | | bcd | bcd | bcd | a |

**Lead**

| Treatment | 0 | 2 | 5 | 9 | 12 |
|---|---|---|---|---|---|
| C | b | a | a | a | a |
| WD | | a | a | a | a |
| S- | a | a | a | a | a |
| WD/S- | | a | a | a | a |

**Sulfur**

| Treatment | 0 | 2 | 5 | 9 | 12 |
|---|---|---|---|---|---|
| C | a | a | ab | b | a |
| WD | | b | b | b | a |
| S- | b | c | c | c | b |
| WD/S- | | c | c | c | b |

**Vanadium**

| Treatment | 0 | 2 | 5 | 9 | 12 |
|---|---|---|---|---|---|
| C | a | a | a | c | ab |
| WD | | a | a | bc | ab |
| S- | a | a | a | c | b |
| WD/S- | | a | a | ab | a |

**Zinc**

| Treatment | 0 | 2 | 5 | 9 | 12 |
|---|---|---|---|---|---|
| C | a | bcd | bcd | cd | b |
| WD | | a | bcd | abc | ab |
| S- | a | cd | cd | d | b |
| WD/S- | | abc | ab | a | a |

**Supplementary Fig. S4**. Effect of treatments on the accumulation of elements in leaves. A, Accumulation profiles of elements over time. Data are means ± S.D of *n* = 4 biological replicates. C: control, WD: water deficit, S-: Sulfur deficiency. Note that day 12 corresponds to three days of rewatering in the WD and WD/S- conditions. B, Letters indicating significant differences between treatments at each time point (days 0, 2, 5, 9, 12). Three separate statistical analyses were performed: comparison of S- and control at day 0 (Student's t-test); comparisons of all treatments from day 2 to day 9 (two-way ANOVA followed by Tukey's HSD tests); comparisons of all treatments at day 12 during the plant recovery (one-way ANOVA followed by Tukey's HSD tests, see methods for details).

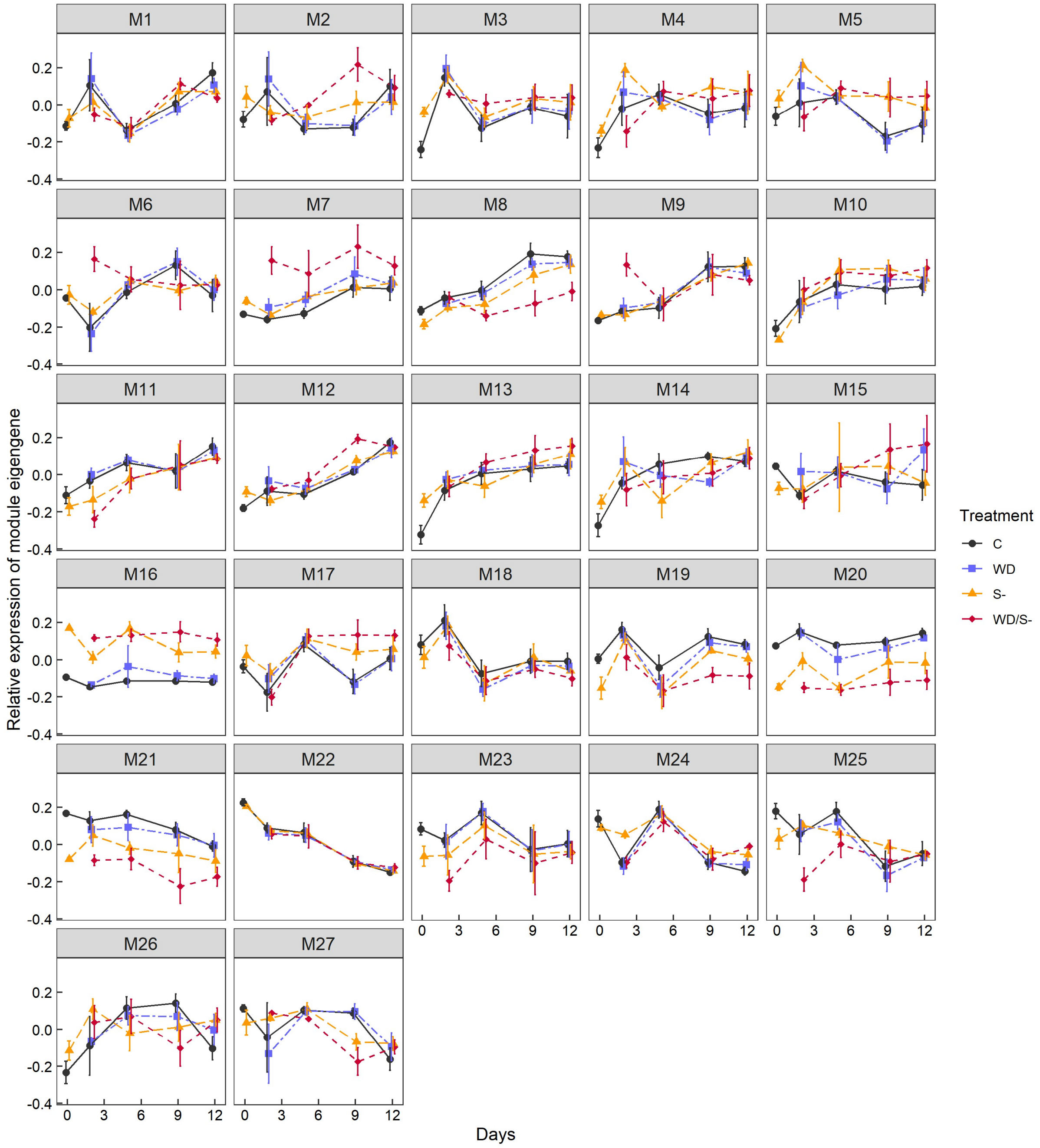


**Supplementary Fig. S5**. Profiles of module eigengenes for modules identified in the co-expression network analysis, performed from transcriptomic data. Data are means ± S.D. for *n* = 4 replicates. C: control, WD: water deficit, S-: Sulfur deficiency. Note that day 12 corresponds to three days of rewatering in the WD and WD/S- conditions.

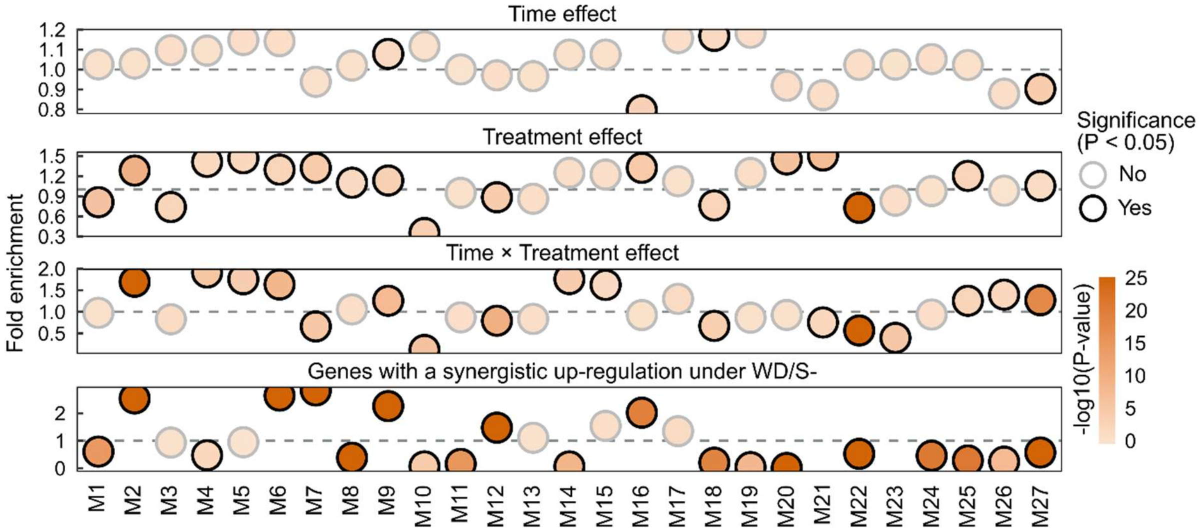


**Supplementary Fig. S6**. Enrichment of specific groups of genes within co-expression modules. Fold enrichment < 1 and > 1 correspond to an under- and over-representation in the module, respectively. Significance (*P* < 0.05, Fisher's exact test) is indicated as bubbles circled in black. Genes with a time, treatment and/or time × treatment interaction effects were identified in Fig. 3A. Genes with a synergistic up-regulation in WD/S- at one or more time points are highlighted in Fig. 6B. The absence of bubbles in certain modules is attributed to missing values (*e.g.* no genes with a synergistic up-regulation under WD/S- were found in module M21 and M23).

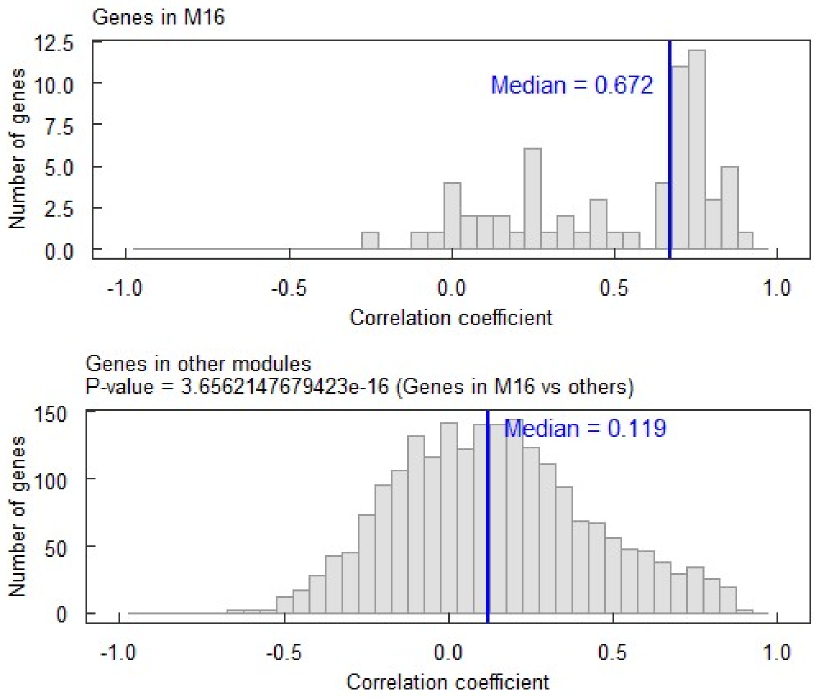


**Supplementary Fig. S7**. Distribution of correlation (Protein vs mRNA, Spearman correlation) coefficients within M16 and other modules. Vertical blue lines indicate medians. The P-value was calculated by comparing correlation coefficients within M16 vs other modules, using a Wilcoxon signed-rank test.

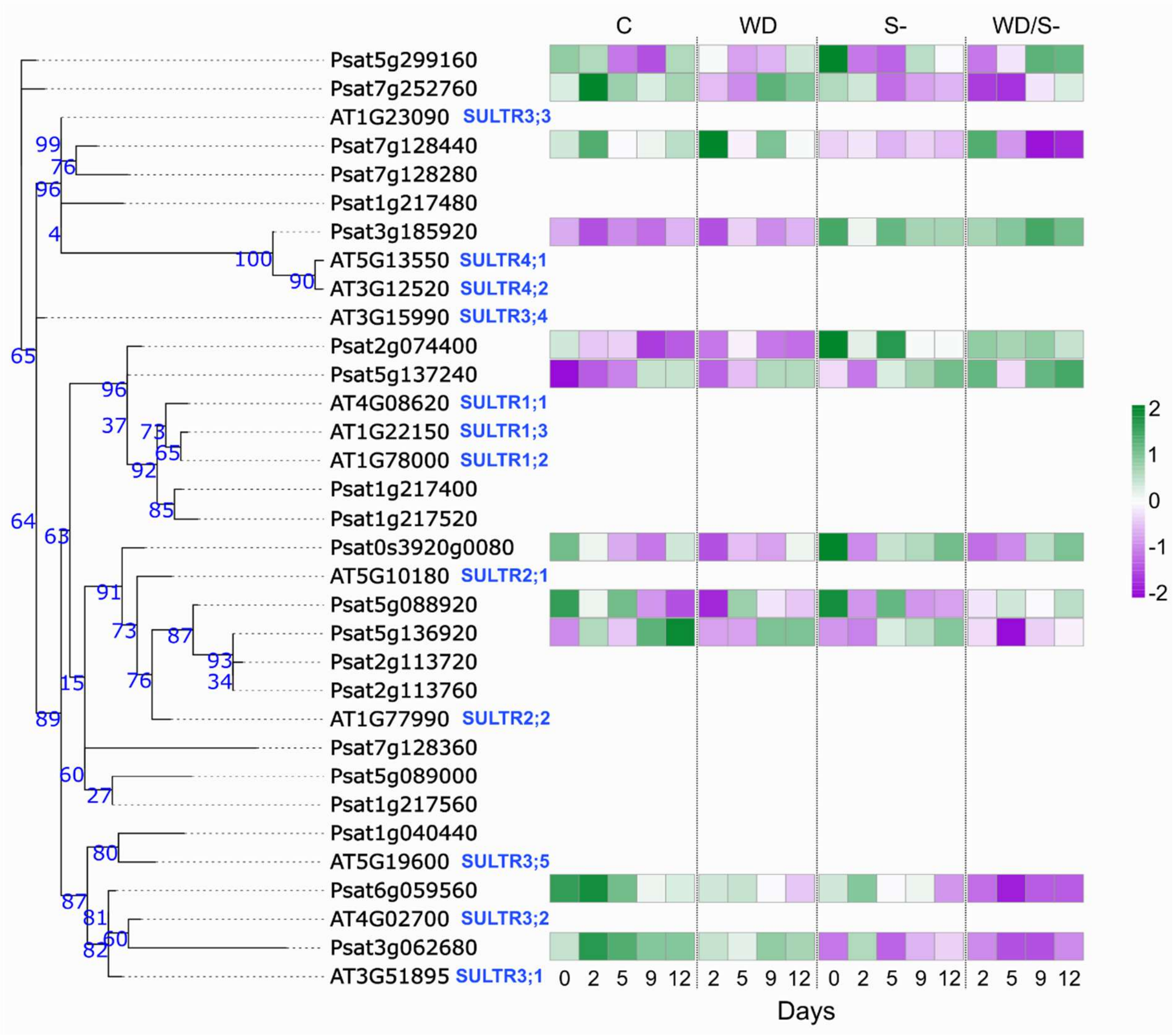


**Supplementary Fig. S8**. Response to stress of sulfate transporters. Phylogenetic tree of sulfate transporters was built using the interactive phylogenetics module of the Dicots PLAZA v5 database (https://bioinformatics.psb.ugent.be/plaza/). Confidence numbers are indicated on the tree branches and provide insights into the reliability of the inferred relationships between sequences. Heatmap represents expression levels of sulfate transporters over time and under control and stress conditions. Data are scaled and are means of *n* = 4 replicates. Only genes identified as expressed in the experiment and selected for the weighted gene co-expression network analysis are represented. Purple and green colors indicate low and high expression levels, respectively. C: control, WD: water deficit, S-: Sulfur deficiency. Note that day 12 corresponds to three days of rewatering in the WD and WD/S- conditions.

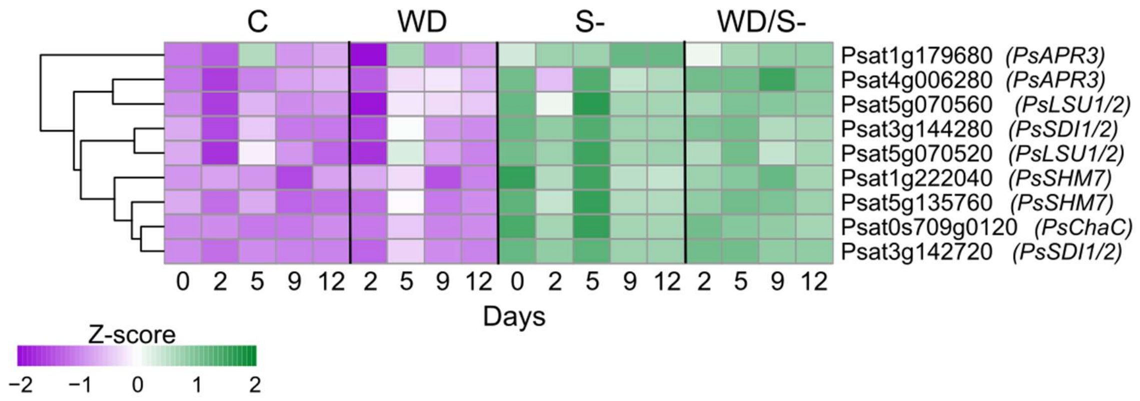


**Supplementary Fig. S9.** Heatmap representing expression levels of OAS cluster genes over time and under control and stress conditions. Best homologous genes of Arabidopsis OAS cluster genes were identified using reciprocal BLASTP. Data are scaled and are means of *n* = 4 replicates. Only genes identified as expressed in the experiment and selected for the weighted gene co-expression network analysis are represented. Purple and green colors indicate low and high expression levels, respectively.

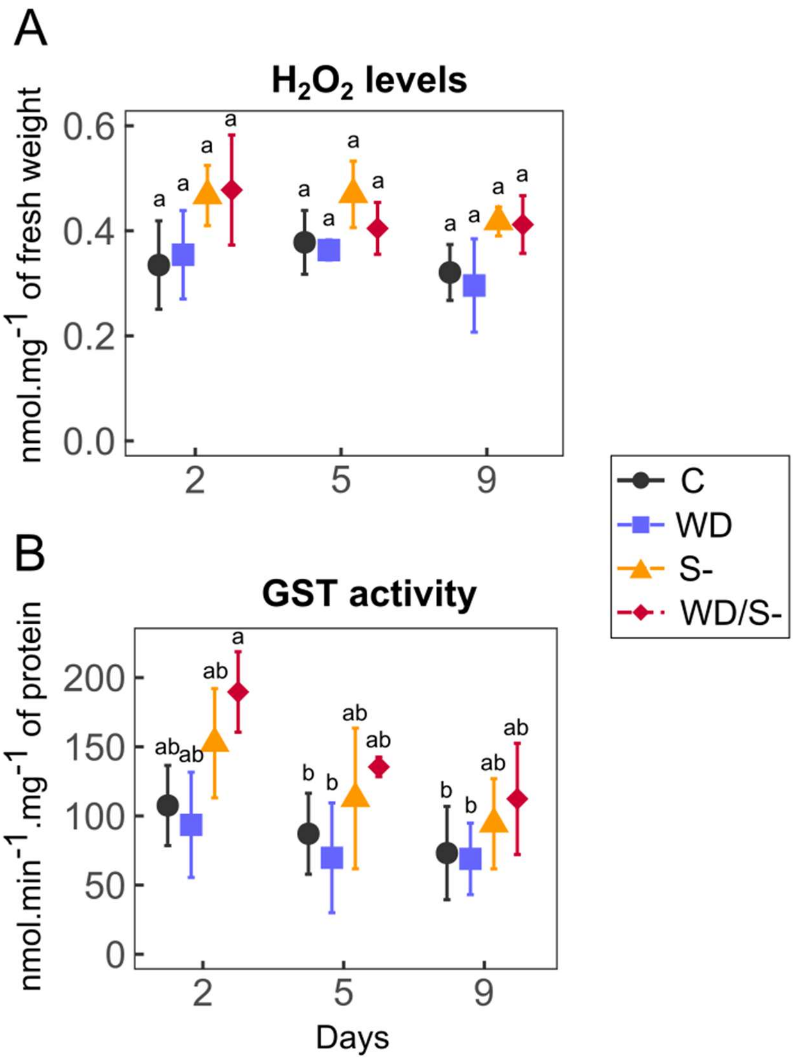


**Supplementary Fig. S10**. Quantification of $H_20_2$ levels (A) and measurement of the glutathione S transferase (GST) activity (B) in leaves at days 2, 5 and 9 (means ± S.D, *n* = 3 replicates). Different letters represent significant differences between treatments.

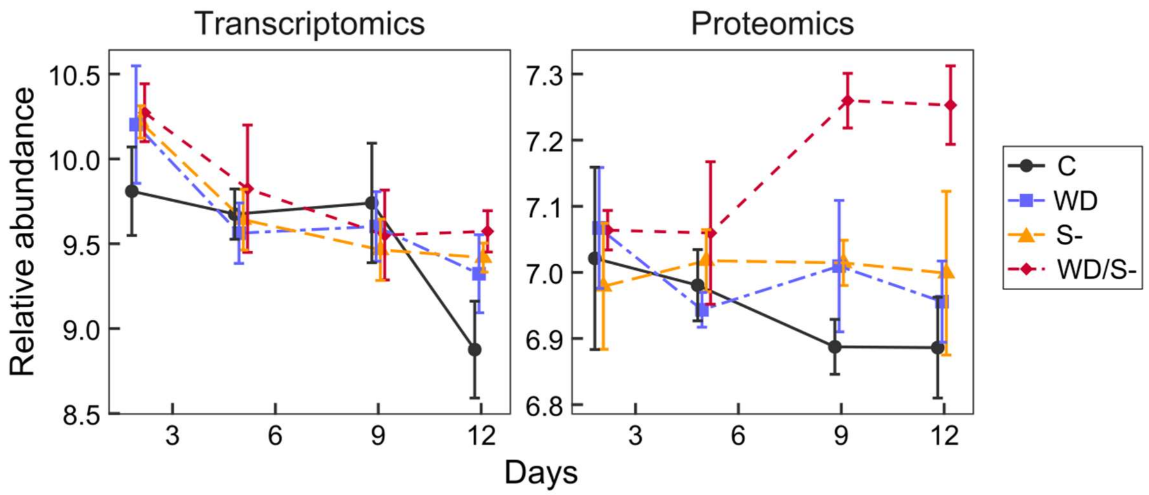


**Supplementary Fig. S11.** Expression profile of *Psat4g008960*, which encodes a Temperature Induced Lipocalin. Data are means ± S.D of $n$ = 4 replicates.
